# GPN-MSA: an alignment-based DNA language model for genome-wide variant effect prediction

**DOI:** 10.1101/2023.10.10.561776

**Authors:** Gonzalo Benegas, Carlos Albors, Alan J. Aw, Chengzhong Ye, Yun S. Song

**Affiliations:** Graduate Group in Computational Biology, University of California, Berkeley; Computer Science Division, University of California, Berkeley; Department of Statistics, University of California, Berkeley; Center for Computational Biology, University of California, Berkeley

## Abstract

Whereas protein language models have demonstrated remarkable efficacy in predicting the effects of missense variants, DNA counterparts have not yet achieved a similar competitive edge for genome-wide variant effect predictions, especially in complex genomes such as that of humans. To address this challenge, we here introduce GPN-MSA, a novel framework for DNA language models that leverages whole-genome sequence alignments across multiple species and takes only a few hours to train. Across several benchmarks on clinical databases (ClinVar, COSMIC, OMIM), experimental functional assays (DMS, DepMap), and population genomic data (gnomAD), our model for the human genome achieves outstanding performance on deleteriousness prediction for both coding and non-coding variants.

## Introduction

With the rising trend in whole-genome sequencing, there is a pressing need to understand the effects of genome-wide variants, which would lay the foundation for precision medicine [1]. In particular, predicting variant deleteriousness is key to rare disease diagnosis [2] and rare variant burden tests [3]. Indeed, a recent review highlights analysis of functional rare variants as the biggest contribution of human genetics to drug discovery [4].

Language models are gaining traction as deleteriousness predictors, with their ability to learn from massive sequence databases and score variants in an unsupervised manner. Given the success of accurately scoring missense variants with protein language models [5–7], it is natural to consider scoring genome-wide variants with DNA language models. For this task, we recently developed the Genomic Pre-trained Network (GPN), a model based on a convolutional neural network trained on unaligned genomes, and showed that it achieves excellent variant effect prediction results in the compact genome of *Arabidopsis thaliana* [8]. The human genome – which harbors a similar number of genes but interspersed over nearly 23 times larger regions and contains much more repetitive elements, most of which may not be functional – is substantially harder to model, however. In fact, previous attempts at unsupervised variant effect prediction with human DNA language models (e.g., Nucleotide Transformer [9]) have shown inferior performance compared to simpler conservation scores. Increasing the scale of the model, data, and compute improves performance, but it can still be poor, even for a model trained for 28 days using 128 top-of-the-line graphics processing units (GPUs) [9].

To address the above challenge, we here introduce GPN-MSA, a novel DNA language model which is designed for genome-wide variant effect prediction and is based on the biologically-motivated integration of a multiple-sequence alignment (MSA) across diverse species using the flexible Transformer architecture [10]. We apply this modeling framework to humans using an MSA of diverse vertebrate genomes [11] and show that it outperforms not only recent DNA language models such as Nucleotide Transformer [9] and HyenaDNA [12] but also current widely-used models such as CADD [13], phyloP [14, 15], ESM-1b [6, 16], Enformer [17], and SpliceAI [18]. Our model took only 3.5 hours to train on 4 NVIDIA A100 GPUs, which is a considerable reduction in the required computing resources compared to the aforementioned Nucleotide Transformer [9]. We anticipate that this massive reduction in computational footprint will enable the efficient exploration of new ideas to train improved DNA language models for genome-wide variant effect prediction.

## Results

GPN-MSA is trained on a whole-genome MSA of 100 vertebrate species (Supplementary Figure 1), after processing (Figure 1a) and filtering (Figure 1b). It is an extension of GPN [8] to learn nucleotide probability distributions conditioned not only on surrounding sequence contexts but also on aligned sequences from related species that provide important information about evolutionary constraints and adaptation (Figure 1c, Methods). It draws inspiration from the MSA Transformer [19], a protein language model trained on MSAs of diverse protein families; it was originally designed for structure prediction but was later shown to achieve excellent missense variant effect prediction performance [5]. Besides the fact that our model operates on whole-genome DNA alignments – which comprise small, fragmented synteny blocks with highly variable levels of conservation, and hence are considerably more complex than protein alignments – there are also essential differences in the architecture and training procedure of GPN-MSA from the MSA Transformer (Methods).

**Figure 1:**
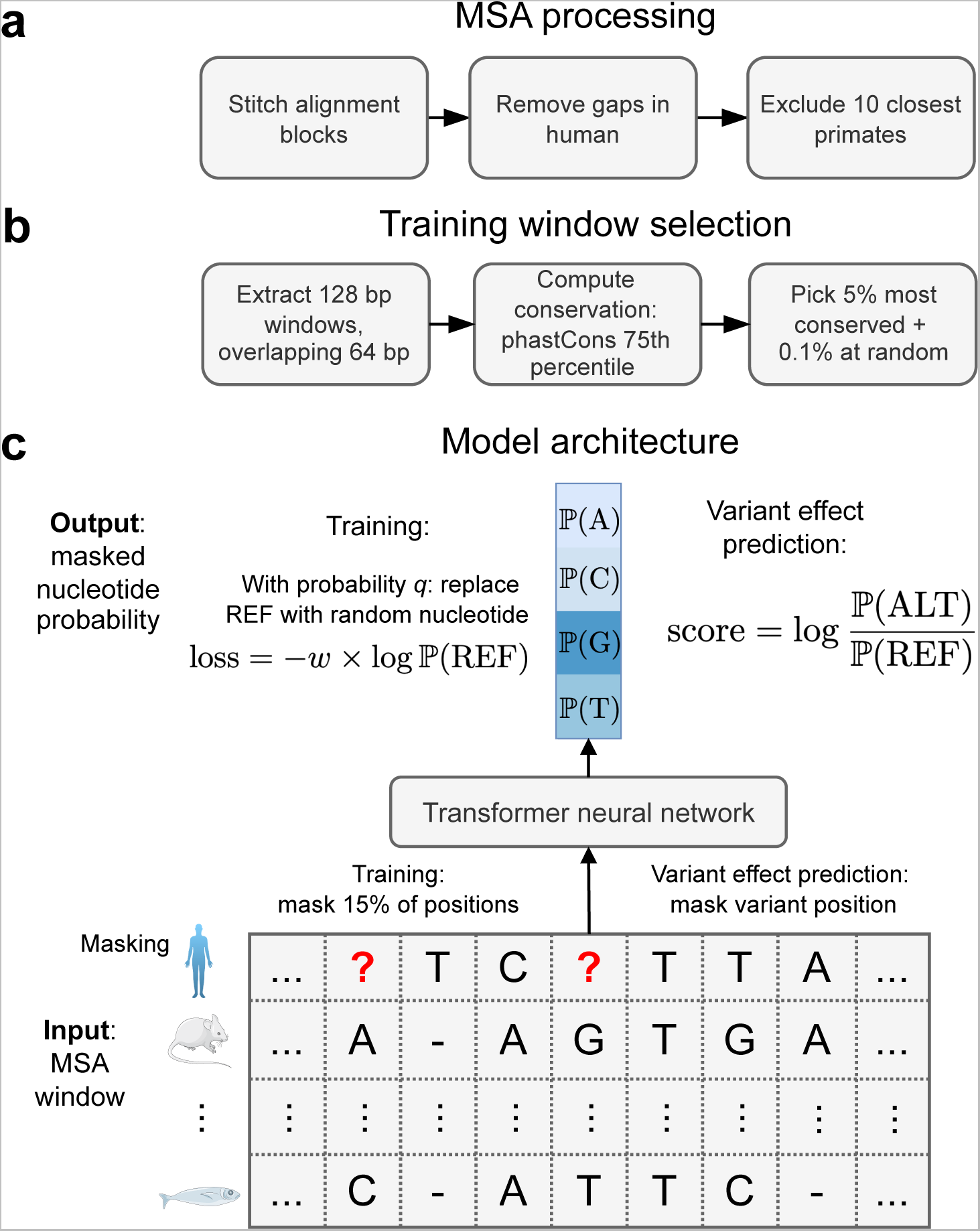
Overview of GPN-MSA. **(a)** MSA processing. Starting with a Multiple Alignment Format file, alignment blocks are stitched together following the order in the human reference. Columns with gaps in the human reference are discarded, followed by the removal of the 10 primate species closest to human (Chimp to squirrel monkey). **(b)** Training window selection. For each 128-bp window along the genome, conservation is computed as the 75^th^ percentile of phastCons. The top 5% conserved windows are chosen alongside a random 0.1% from the remaining windows. **(c)** Model architecture. The input is a 128-bp MSA window where certain positions in the human reference have been masked, and the goal is to predict the nucleotides at the masked positions, given the context across both columns (positions) and rows (species) of the MSA. During training, 15% of the positions are masked. During variant effect prediction, only the variant position is masked. The sequence of MSA columns is processed through a Transformer neural network resulting in a high-dimensional contextual embedding of each position. Then, a final layer outputs four nucleotide probabilities at each masked position. The model is trained with a weighted cross-entropy loss, designed to downweight repetitive elements and up-weight conserved elements (Methods). As data augmentation in non-conserved regions, prior to computing the loss, the reference is sometimes replaced by a random nucleotide (Methods). The GPN-MSA variant effect prediction score is defined as the log-likelihood ratio between the alternate and reference allele. REF: reference allele. ALT: alternate allele. Mouse and fish icons are from Servier (https://smart.servier.com/).

By utilizing the MSA as auxiliary information, GPN-MSA can accurately predict nucleotides from their context, especially in functional regions (Supplementary Table 1). At sites where the reference allele differs from the inferred ancestral allele [20], GPN-MSA usually favors the ancestral allele (Supplementary Table 2). However, predicting the human reference is just a pretext task. What we really care about is the likelihood assigned to human genetic variants that have not been seen during training. Conservation statistics computed on an MSA column, from simple frequencies to more complex phylogeny-aware *p*-values [14], are intuitive and powerful measures of deleteriousness. GPN-MSA is designed to process conservation information across multiple MSA columns, as has been exploited by earlier models such as phastCons [21] based on a hidden Markov model. To illustrate GPN-MSA’s power beyond single-column statistics and its ability to leverage genomic context, we note that, even at perfectly conserved positions, GPN-MSA assigns more deleterious scores to loss-of-function (e.g., stop gain/loss, splice donor/acceptor variant) and missense variants compared to synonymous variants (Figure 2a). Furthermore, variants with extreme GPN-MSA log-likelihood ratios tend to have lower minor allele frequencies (MAF) than variants with extreme log-likelihood ratios based on MSA column frequencies, suggesting that GPN-MSA is a better estimator of deleteriousness (Figure 2b).

**Figure 2:**
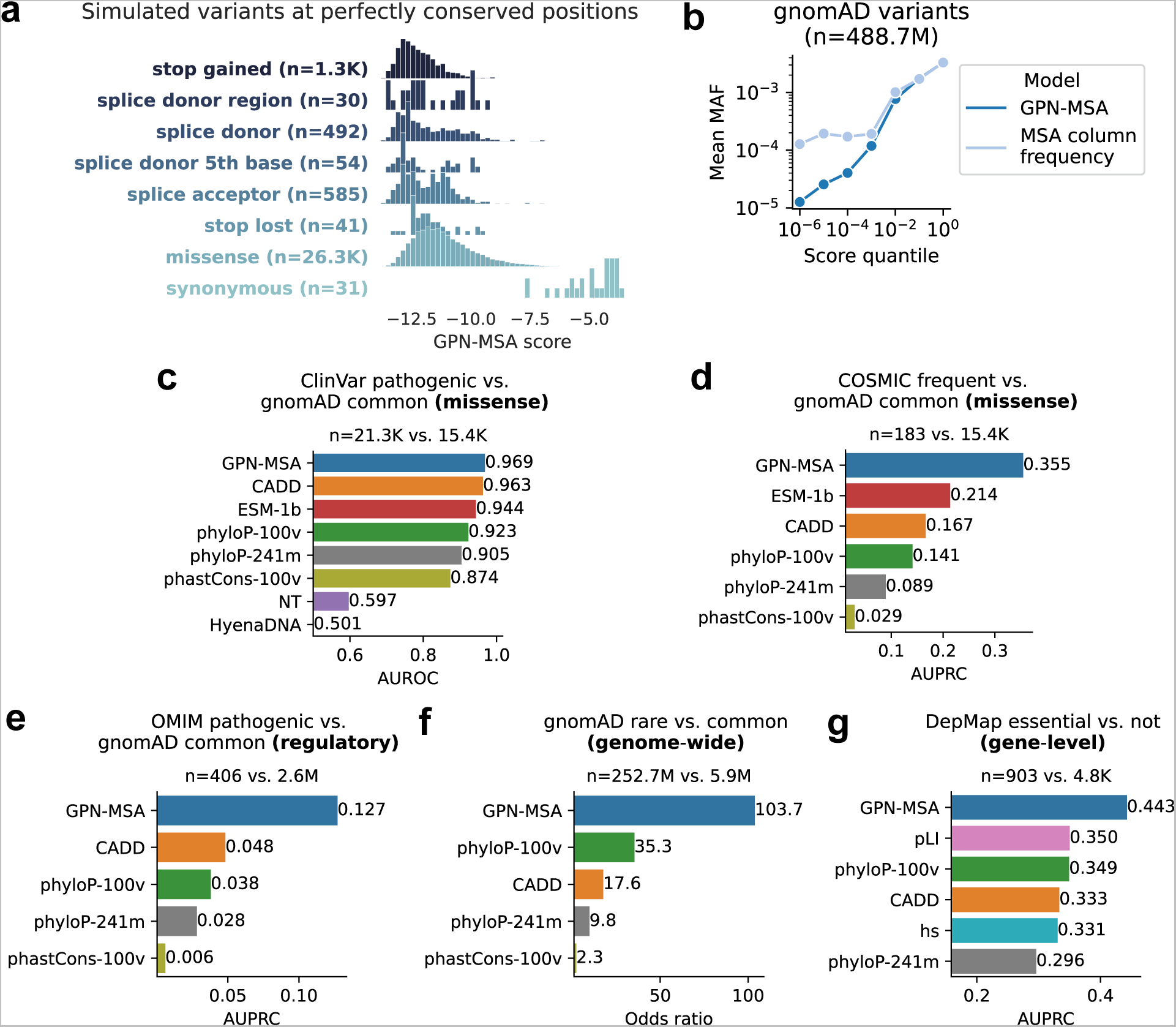
Variant effect prediction results. **(a)** Distribution of GPN-MSA scores at positions in held-out chromosome 22 where the model sees perfect nucleotide identity in the MSA column (i.e. perfect identity in the 89 non-human species), across categories. **(b)** Mean minor allele frequency (MAF) for different score quantile bins ([0, 10*^−^*^6^), (10*^−^*^6^, 10*^−^*^5^]*, …,* (10*^−^*^1^, 0]) in the full set of gnomAD bi-allelic sites. The MSA column frequency score is the log-likelihood ratio based on the empirical column frequencies, with a pseudocount of 1. To break ties, we added a very small random number to each score (the pattern across random seeds was stable). **(c)** Classification of ClinVar pathogenic vs. gnomAD common missense variants. NT: Nucleotide Transformer. **(d)** Classification of COSMIC frequent (frequency *>* 0.1%) vs. gnomAD common missense variants. **(e)** Classification of OMIM pathogenic vs. gnomAD common regulatory variants. We matched OMIM promoter variants with gnomAD upstream-of-gene variants, enhancer with intergenic, and “all” with the union of the matches of the specific categories. **(f)** Enrichment of rare (singletons) vs. common (MAF *>* 5%) gnomAD variants (subset) in the tail of deleterious scores (the threshold was chosen such that each score makes 30 false discoveries). Odds ratios and *p*-values were computed using one-sided Fisher’s exact test. All shown odds ratios have *p*-value *<* 0.05. **(g)** Classification of DepMap essential vs. non-essential genes based on the 1^st^ percentile of scores across the gene. Essential genes are defined as those *>* 1000 DepMap cell lines are dependent on and non-essential genes are defined as those no cell line is dependent on. AUROC: area under the receiving operating characteristic curve. AUPRC: area under the precision-recall curve. MAF: minor allele frequency.

We demonstrate the capability of GPN-MSA to improve unsupervised deleteriousness prediction on several human variant datasets (Methods). We emphasize that only the reference genome is used to train GPN-MSA and that no human variant dataset is utilized in training. Nevertheless, GPN-MSA can still capture several functional attributes of variants, such as epigenetic marks and the impact of natural selection (Supplementary Figure 2).

For evaluation, we first consider the classification of ClinVar [22] pathogenic vs. common missense variants in gnomAD [23]. GPN-MSA substantially outperforms other human DNA language models such as Nucleotide Transformer (NT) [9], with the largest amount of parameters (2.5 B), as well as HyenaDNA [12], with the largest context size of 1 Mb (Figure 2c, Supplementary Figure 3a). We also find that GPN-MSA achieves improved performance compared to genome-wide predictors CADD [24] and phyloP [14, 15], as well as the missense-specific ESM-1b [6, 16]. These results are based on using common variants as controls instead of ClinVar benign-labeled variants, as recommended by the developers of CADD to reduce ascertainment bias [13]. When using benign-labeled variants in ClinVar as controls, GPN-MSA performs slightly behind CADD and ESM-1b (Supplementary Figure 4). A type of bias that is particularly problematic for benchmarking computational predictors is that a variant with a high population frequency can be labeled as benign (ACMG/AMP classification criterion BA1), but this label can be removed if computational methods predict it to be pathogenic (criterion PP3) [25]. Therefore, popular predictors among ClinVar submitters (such as older predictors), or newer predictors with high similarity to the latter, will have an unfair advantage.

Next, we consider the classification of somatic missense variants frequently observed across cancer tumors (COSMIC, the Catalogue of Somatic Mutations in Cancer [26]) vs. gnomAD common missense variants. Because of the extreme class imbalance in this case, we focus on the precision and recall metrics. GPN-MSA again achieves the highest performance, with substantial margins of improvement over other models (Figure 2d, Supplementary Figure 3b).

We also evaluate on deep mutational scanning (DMS) experimental data [27] for 31 human proteins (Supplementary Table 3). GPN-MSA and CADD perform comparably on classifying variants labeled according to protein-specific binarization (Methods), with the former slightly outperforming the latter on the AUPRC metric (Supplementary Figure 5). They both compare favorably with phyloP and phastCons. However, the protein language model ESM-1b achieves the best overall performance on this task, which is not surprising, as these data are not well suited for genome-wide variant effect predictors, given the lack of introns and other genomic contexts. Protein language models therefore have the advantage in this specific setting given that they are trained on a much more diverse set of protein sequences. Nevertheless, GPN-MSA performs better than ESM-1b on some proteins (e.g., TADBP, for which GPN-MSA AUROC is 0.83, while ESM-1b AUROC is 0.75). Exploring the conditions under which one model outperforms another, and integrating the strengths of both DNA and protein language models, presents an intriguing avenue for future research. As another note of caution, none of the models performs exceedingly well relative to ClinVar results.

Moving on to regulatory variants, we evaluate on the classification of a curated set of variants implicated in Mendelian disorders (OMIM, Online Mendelian Inheritance in Man [28]) vs. gnomAD common variants. We again consider precision and recall because of the extreme class imbalance, and find that GPN-MSA achieves the best performance overall, as well as in each variant category (Figure 2e, Supplementary Figure 6, Supplementary Figure 3c). NT exhibits poor performance compared to other models (Supplementary Figure 6). For several variant categories, CADD’s precision increases from near zero as recall increases, which indicates that a substantial fraction of its top discoveries are actually false (Supplementary Figure 3c). One example of a deleterious variant that was assigned an extreme score by GPN-MSA is rs606231231, lying in the well-known ZRS enhancer that controls the expression of *SHH* at the long range of 1 Mb (Supplementary Figure 7). This variant is associated with polydactyly [29] and has been experimentally verified to alter gene expression in mouse limb [30]. Another example is rs1367115848, disrupting HNF4 binding at the *F7* promoter and causing severe factor VII deficiency [31] (Supplementary Figure 8).

Following this, we further evaluate on the enrichment of rare vs. common gnomAD variants in the tail of the distribution of deleteriousness scores. Deleterious variants should be under purifying selection and hence their frequencies in populations should tend to be lower. Therefore, if a variant effect predictor is accurate, we expect rare variants to be enriched compared to common variants for extreme deleteriousness scores. GPN-MSA achieves the highest enrichment overall (Figure 2f), as well as within most variant categories, with different margins (Supplementary Figure 9, Supplementary Figure 10). In the case of intronic variants, it also outperforms SpliceAI [18], a state-of-the-art splicing predictor. There is one category where GPN-MSA performs behind CADD: splice-region variants (here we group variants immediately close to the exon borders, such as splice donors and acceptors). To be clear, GPN-MSA generally assigns extreme scores to these variants (Supplementary Figure 11a); the challenge is understanding which variants are *not* deleterious and more likely to be common in the population. Many of the features utilized by CADD—such as gene and transcript identifiers—could help in this task and potentially be integrated into GPN-MSA to improve performance. We note that the overall genome-wide performance in Figure 2f is not merely an averaging of the performances in the different categories; it also involves scoring variants relative to each other across these categories. On a separate enrichment analysis of low-frequency vs. common gnomAD variants in non-exonic regions, GPN-MSA achieves a substantially improved performance over Enformer [17] (Supplementary Figure 12, Supplementary Figure 13).

GPN-MSA also outperforms other methods when subsetting to putatively conserved, neutral, or accelerated positions in the genome (Supplementary Figure 14). While we observe that SpliceAI and Enformer, which are functional genomics models, perform worse than the simpler phyloP in deleteriousness prediction, we note that this is an application they were not designed for. It is also worth noting that although phyloP-241m (fit to the 241-way Zoonomia mammalian alignment) was recently proposed as a deleteriousness predictor [15], the older vertebrate phyloP-100v actually achieves better results in many of our benchmarks.

Additionally, we evaluate our model on the classification of essential genes using the DepMap cancer dependency data [32]. This dataset contains gene essentiality measurements based on genome-scale CRISPR knockout screens on over 1000 cancer cell lines. Essential genes are supposed to be more intolerant to deleterious mutations and consequently, their variants tend to have overall higher impacts. We summarize variant effect predictions by taking the tail of deleterious scores across each gene and investigate how well each method can classify essential versus non-essential genes among the cell lines. GPN-MSA outperforms other genome-wide variant effect predictors, as well as methods (pLI [33] and *hs* [34]) specifically designed for gene essentiality prediction (Figure 2g).

To understand the importance of different components of our model, we perform an ablation study and assess the impact on variant effect prediction performance. We here summarize the main takeaways (Supplementary Table 4). Firstly, the inclusion of the MSA is critical for GPN-MSA’s high performance. Secondly, the simple MSA column frequency is already a good predictor for coding variants, but does a poor job with non-coding variants. Thirdly, in our case training on conserved regions is better than the usual approach of training on the whole genome. Lastly, with an MSA of diverse vertebrate genomes, we see diminishing returns of increased window size for deleteriousness prediction. More details are provided in Methods.

We provide pre-computed scores for all *∼*9 billion possible single nucleotide variants in the human genome, as well as sequence logos [35] in the UCSC Genome Browser [36, 37] (example in Supplementary Figure 15). Examining the complete gnomAD variant set, there seems to be a near-linear relationship between the GPN-MSA score bin and the logarithm of the average minor allele frequency within that specific bin (Supplementary Figure 16). We believe that the deleteriousness of GPN-MSA scores should be interpreted as a continuum; if a hard threshold is helpful, we recommend a cutoff around *−*7, based on the distribution of scores in different datasets (Supplementary Figure 17). Incidentally, the bimodality of score distribution for frequent variants in COSMIC suggests that many of them could be passenger mutations (Supplementary Figure 17b). Additionally, we recommend using scores only to compare variants relative to each other, and warn against interpreting them as calibrated fitness estimates (Methods).

## Discussion

To recapitulate, our main contributions are threefold. First, we propose the first DNA language model operating directly on a whole-genome alignment. Second, we demonstrate outstanding performance in humans on a number of clinically-relevant variant effect prediction datasets. While there already exist many effective missense variant effect predictors, we anticipate that our ideas, and readily available GPN-MSA scores, will be particularly helpful to interpret variation in non-coding regions. Lastly, the general approach we have developed for humans is computationally efficient, which would enable future research in the field.

In the rapidly advancing landscape of DNA language modeling, scaling up model and context sizes have been the primary avenues of exploration [9, 12, 38]. In contrast, our present work focuses on the explicit modeling of related sequences (known as retrieval augmentation in natural language processing [39]). This has led to a highly computationally efficient model and state-of-the-art variant effect prediction performance for genome-wide variants. It remains to be explored how useful GPN-MSA’s learned representations would be for downstream applications, e.g., for genome annotation or gene expression prediction. Expanding the context length, possibly through leveraging recent technical developments [12], might be beneficial for such tasks.

Our modeling approach also differs from earlier works, in that we train mostly on conserved regions of the genome to increase the proportion of functional as opposed to neutral sites. This strategy may potentially miss some functional sites, including those in primate or human-specific regions, fast-evolving regions, or in hard-to-align regions. We also recognize the challenge in balancing data quality (i.e., sequence information content) with data quantity during training, given that larger models have a higher risk of overfitting when trained on smaller amounts of data. We believe that these are critical considerations for researchers seeking to adopt our present strategy. The masked language modeling objective can be too easy if sequences very similar to the human genome are included in the MSA, resulting in the learned probability distribution being not very useful for variant effect prediction. This observation has led us to exclude most primate genomes during training. To tackle this limitation, we are actively exploring alternative training objectives which are aware of phylogenetic relationships. We are also exploring how best to integrate population genetic variation information, instead of relying on a single reference genome.

In our view, one of the most promising applications of GPN-MSA is effective genome-wide rare variant burden testing, which has been mostly restricted to coding regions [40]. We envision that several other statistical genetics tasks can be empowered by GPN-MSA, such as functionally informed fine-mapping [41]; polygenic risk scores [42] and related variant randomness tests [43]; and variant prioritization in integrative analyses.

Sequence models (such as phyloP and GPN-MSA) might achieve better deleteriousness prediction results but are still less interpretable than functional genomics models such as SpliceAI and Enformer. While both functional genomics models and DNA language models have much room for independent improvement, it is likely that jointly modeling DNA sequence and functional genomics may have the biggest impact.

## Methods

### MSA Processing

The multiz [46] whole-genome alignment of 100 vertebrates was downloaded from https://hgdownload.soe.ucsc.edu/goldenPath/hg38/multiz100way/maf/. Contiguous alignment blocks were stitched together using the multiz utility maf2fasta and any columns with gaps in human were removed. The 10 primate species closest to human were removed. We also downloaded associated conservation scores phastCons [47] https://hgdownload.soe.ucsc.edu/goldenPath/hg38/phastCons100way and phyloP [14] https://hgdownload.soe.ucsc.edu/goldenPath/hg38/ phyloP100way.

### Training Region Selection

Instead of training on the whole genome, we focused on the most conserved genomic windows, aiming to emphasize functionally-important regions such as exons, promoters and enhancers. The conservation of a genomic window was defined as the 75^th^ percentile of phastCons scores in the window. We then chose a cutoff; in our current experiments we included the top 5% most conserved windows. We also included 0.1% of the remaining windows of the genome to ensure there is no extreme distribution shift when performing variant effect prediction in non-conserved regions. The reverse complement of each selected window was added as data augmentation. Chromosome 21 was held out for validation (early-stopping) and chromosome 22 was held out for evaluating language modeling performance metrics such as perplexity. We used all chromosomes for evaluating the separate task of variant effect prediction.

### Model Architecture

We adopt the general approach of masked language modeling [48]. As a general caveat, in this work we did not systematically tune hyperparameters, so they are likely far from optimal. The input is a 128-bp MSA window where certain positions in the human reference have been masked, and the goal is to predict the nucleotides at the masked positions, given its context across both columns (positions) and rows (species) of the MSA. During training, 15% of the positions are masked. During variant effect prediction, only the variant position is masked. The one-hot encodings of nucleotides from different species at each position are first concatenated. Then, the sequence of MSA columns is processed through a Transformer neural network (RoFormer [49]) resulting in a high-dimensional contextual embedding of each position. Then, a final layer outputs four nucleotide probabilities at each masked position. The model is trained on the reference sequence with a weighted cross-entropy loss.

We downweight repeats and upweight conserved elements so that incorrect predictions in neutral regions are penalized less. We introduce a smoothed version of phastCons, phastCons*_M_*, as the max of phastCons over a window of 7 nucleotides. The goal was to not only give importance to conserved regions, but to regions immediately next to them. The loss weight *w* is defined as follows:

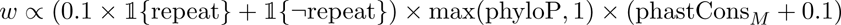

which includes 10-fold downweighting on repetitive elements [8] plus upweighting based on both phyloP and phastCons*_M_*.

As data augmentation in non-conserved regions, prior to computing the loss, the reference is replaced by a random nucleotide with a certain probability *q*:

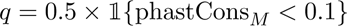

As a result, the probability of keeping the reference nucleotide is 0.5+ 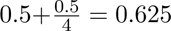 and the probability of replacing it with each of the alternative nucleotides is 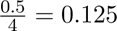. The intention is to guide the model to assign more neutral scores in non-conserved regions.

Our code is based on the Hugging Face Transformers library [50]. All models were trained with default hyperparameters ^1^ (e.g. 12 layers with 12 attention heads each) except for the ones listed in Supplementary Table 5. The total number of parameters is approximately 86 million. We performed early stopping based on validation loss. We trained the model for a max of 30 thousand steps (*≈* 14 epochs) in approximately 3.5 hours using 4 NVIDIA A100 GPUs.

The GPN-MSA variant effect prediction score is defined as the log-likelihood ratio between the alternate and reference allele. In our experiments, we average the predictions from the positive and negative strands. With our 4 NVIDIA A100 GPUs, we manage to score approximately 25 million variants per hour.

### Differences between GPN-MSA and MSA Transformer

While the MSA Transformer takes as input an arbitrary set of aligned sequences, GPN-MSA is trained on sequences from a fixed set of species. This allows simpler modeling of the MSA as a sequence of fixed-size alignment columns, reducing computation and memory requirements. Variant effect prediction, masking only the target sequence (in our case, human), is identical [5]. Since variant effect prediction is our main goal, during training we also only mask positions from the target sequence. The MSA Transformer, however, proposes masking MSA entries at random during training, based on results from structure prediction, their intended application.

### Language modeling metrics

The median perplexity and accuracy were computed for the test chromosome (chr 22). Perplexity is defined as the exponentiation of the cross-entropy loss, which is equivalent to 1 over the probability given to the correct nucleotide. The ancestral alleles [20] were downloaded from https://ftp.ensembl.org/pub/release-109/fasta/ancestral_alleles/. The accuracy was computed for test chromosome positions where the ancestral and reference alleles differ.

### Ablation Study

We performed an ablation study to understand the impact of each of our design choices on variant effect prediction when modified independently (Supplementary Table 4). For each setting, three replicate models with different seeds were trained, where applicable. Since we hold the rest of the hyperparameters fixed, results should be interpreted as differences given a similar training procedure and compute budget. There are many crucial hyperparameters worth investigating further. These include the size of the model, the number of iterations, and the learning rate schedule, all of which would most certainly affect the performance of each ablated model.

While varying window size has been mainly motivated by the question of how much context can be leveraged by the model to make a prediction, it also affects two other components of the training workflow. Firstly, it affects which positions of the genome are used for training, given that we first split the genome into windows of a given size and filter them according to the 75^th^ phastCons percentile — this statistic will be biased towards certain regions over others according to the window size. Secondly, it will influence the number of data points (windows) or the total number of tokens (base pairs) used for training, which can affect optimization and generalization. With these considerations in mind, to study how much the model uses context, we decided to reduce the window size at variant effect prediction time rather than retrain from scratch. For the increased window size we did retrain the model from scratch, but for better performance we encourage future studies to also vary other hyperparameters, particularly increasing the percentage of the genome used for training, to reduce the chance of overfitting.

Details on the ablations:

- w/o MSA: the model is only trained on the human sequence, without access to other species.
- MSA frequency: variants are scored using the log-likelihood ratio of observed frequencies in the MSA column, with a pseudocount of 1.
- Combined phyloP and phastCons: the same combination used to upweight the loss.
- Train on 50%-100% most conserved: expand the training region from the smaller 5% most conserved to a larger set with less overall conservation.
- Include closest primates: do not filter out from the MSA the 10 primates closest to human.
- 51 mammals: subset the vertebrate MSA to only mammals (51, besides human).
- 51 vertebrates: subset the vertebrate MSA from 89 to 51 vertebrates (besides human). The subset is made at random except for the closest species (bushbaby) which is deterministically chosen.
- Don’t upweight conserved: do not upweight the loss function on conserved elements.
- Don’t replace non-conserved: do not replace the reference in non-conserved positions with random nucleotides when computing the loss function.
- Increased window size (256): the whole training procedure (including window selection) is repeated with window size 256.
- Reduced window size (4-64): the default model is shown a smaller window at variant effect prediction time.

We now describe the results. While all metrics can distinguish large differences in performance, the ClinVar and gnomAD metrics are based on larger sample sizes and, therefore, should be more adequate for investigating more subtle differences. Modeling the single human sequence instead of the MSA has by far the biggest impact. We note that alignment-based methods, including simple ones such as MSA frequency, usually achieve a high performance in metrics for missense variants (ClinVar and COSMIC) but not for genome-wide variants (OMIM and gnomAD). Including primate species close to human, or training on less conserved regions, also have a large impact on performance. Of relatively minor impact are subsetting to 51 mammals or 51 vertebrates, removing the upweighting of conserved elements or removing the data augmentation procedure of replacing nucleotides in non-conserved positions. Increasing the window size provided at variant effect prediction from 4 to 128 shows diminishing returns. A model trained with the doubled window size of 256 actually shows worse gnomAD mean performance across random seeds, but a comparable max performance. This instability across runs could potentially be reduced by increasing the percentage of the genome used for training.

### Variant Effect Prediction (VEP) glossary

Variant consequences were obtained by running Ensemble VEP Release 109 [44] with arguments --most severe --distance 1000.

For *in silico* mutagenesis analysis, we analyzed the scores for a random subset of 10M simulated variants in test chromosome 22. Our data augmentation procedure of replacing the reference with another nucleotide in non-conserved regions causes an artificial shift in the distribution of scores (Supplementary Figure 11). The peak observed in the distribution of scores (Supplementary Figure 11a) roughly coincides with 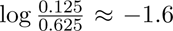, related to the probability that the alternate and reference alleles have been exchanged as data augmentation in non-conserved regions. More importantly, and regardless of the previous point, we consider likelihood ratios to be systematically lower than the ratios expected based on differences in fitness. To illustrate this point, we note that the distribution of GPN-MSA scores for synonymous variants is centered around *−*3, meaning that the alternate allele is usually 20 times less likely than the reference allele, while we would expect them to be equally likely (corresponding to a score of 0, as observed in GPN [8]). The current training procedure still favors assigning a higher probability to the most frequent allele in the MSA even when the fitness is uniform, an issue which may be mitigated by explicitly modeling phylogenetic relationships. Nevertheless, it is clear from the benchmark results that the scores, used in a relative fashion, are very powerful at discriminating deleterious from neutral variants.

We also analyzed the scores at all positions in chromosome 22 where the aligned species (regardless of human, which is masked) have the same nucleotide.

We summarize datasets and their provenance, metrics used to evaluate each dataset, and technical details in constructing VEP scores below.

#### VEP data sources

- ClinVar [22]: downloaded release 20230730.
- COSMIC [26]: downloaded Cosmic MutantCensus v98 GRCh38.tsv.gz and computed frequency as the proportion of samples containing the mutation, restricting to whole-genome or whole-exome samples.
- OMIM [28]: downloaded a set of curated pathogenic regulatory variants.
- gnomAD [23]: downloaded version 3.1.2 and filtered to variants with allele number of at least 25 000, besides the official quality-control flags. We defined common variants as those with minor allele frequency (MAF) *>* 5% and rare variants as the singletons.
- DMS [27]: downloaded the human protein data in ProteinGym v0.1 (Supplementary Table 3). For genomic variant effect predictors, we took the most deleterious score among all the SNVs causing the aminoacid substitution reported in ProteinGym.
- DepMap [32]: downloaded the gene dependency data CRISPRGeneDependency.csv in the Public 23Q4 release. The data were further summarized into per-gene essentiality by counting the number of dependent cell lines. A cell line is defined as dependent on the gene if the probability of dependency is greater than 0.5.

#### Main VEP metrics

- ClinVar: area under the receiving operating characteristic curve (AUROC) for classification of ClinVar “Pathogenic” vs. gnomAD common missense variants.
- COSMIC: area under the precision-recall curve (AUPRC) for classifying COSMIC frequent (frequency *>* 0.1%) vs. gnomAD common missense variants.
- OMIM: AUPRC for classification of OMIM pathogenic vs. gnomAD common regulatory variants. We matched OMIM promoter variants with gnomAD upstream-of-gene variants, enhancer with intergenic, and “all” with the union of the matches of the specific categories.
- gnomAD: enrichment of rare vs. common gnomAD variants in the tail of deleterious scores (the threshold defined by allowing a small number of false discoveries, such as 30). We grouped certain variant consequences with small samples sizes, as follows. “splice-region” groups Ensembl categories splice donor, splice acceptor, splice donor 5th base, splice donor region, splice region and splice polypyrimidine tract. “start-or-stop” groups Ensembl categories start lost, stop gained or stop lost.
- DepMap: AUPRC for classifying essential genes (number of dependent cell lines *>* 1000) vs. non-essential genes (number of dependent cell lines is 0) among DepMap cancer cell lines. To obtain gene essentiality estimates from VEP scores, we first took the mean prediction at each genomic position across the three alternative alleles. Then we summarized across positions by extracting the 1st percentile score, indicative of the most deleterious impact within the gene. All positions within the one selected Ensembl transcript in gnomAD v2 Constraint table were considered for each gene. We included genes where all the benchmarked methods have available predictions at all positions.

#### Additional VEP metrics

- gnomAD (Enformer set): enrichment of low-frequency (0.5% *<* AF *<* 5%) vs. common (MAF *>* 5%) gnomAD non-exonic variants. We use low-frequency instead of rare because of lack of Enformer pre-computed scores.
- ClinVar pathogenic vs. benign: AUROC for classification of ClinVar “Pathogenic” vs. ClinVar “Benign” missense variants.
- DMS: AUROC and AUPRC for binary classification where labels were defined by the protein-specific binarization cutoff provided in ProteinGym, except for TADBP. All models do not perform well on predicting the relative toxicity measurement for TADBP variants [45], but they work much better on predicting the absolute value of relative toxicity. We used a cutoff of 0.125 on the absolute value of relative toxicity to binarize the DMS data for TADBP.
- gnomAD (ablation set): enrichment of rare vs. common gnomAD variants in the tail of deleterious scores. Because of the large size of the full data, we subset the rare variants to match the number of common variants per consequence.

#### VEP scores

- GPN-MSA: log-likelihood ratio between alternate and reference allele. Predictions from both strands were averaged.
- CADD: raw scores (v1.6), negated so lower means more deleterious.
- phyloP-100v: computed on 100 vertebrates alignment, negated so lower means more deleterious.
- phastCons-100v: computed on 100 vertebrates alignment, negated so lower means more deleterious.
- phyloP-241m: computed on 241 mammals alignment, negated so lower means more deleterious.
- Nucleotide Transformer (NT): the most powerful version (2.5b-multi-species), with 2.5B parameters and trained on 850 species. The center 6-mer was masked and the score was computed as the log-likelihood ratio between alternate and reference 6-mer. Predictions from both strands were averaged. Given the high computational requirements, we only scored variants for ClinVar and a subset of OMIM variants.
- HyenaDNA: the most powerful version (large-1m-seqlen-hf), with 54.6M parameters and a context length of 1 Mb. The log-likelihood ratio between the alternate and reference sequence was computed. Predictions from both strands were averaged. Given the very high computational requirements, we only scored variants for ClinVar.
- ESM-1b: precomputed log-likelihood ratios between alternate and reference alleles were obtained in protein coordinates [6]. For variants affecting multiple isoforms, the minimum (most deleterious) score was considered.
- SpliceAI: precomputed scores recommended for variant effect prediction (spliceai scor es.masked.snv.hg38.vcf.gz) were downloaded from https://basespace.illumina.com/ s/otSPW8hnhaZR. The authors do not recommend any specific way of computing a single deleteriousness score. We scored variants using minus the maximum absolute delta in splice acceptor or donor probability in any gene.
- Enformer: precomputed scores for variants with minor allele frequency (MAF) greater than 0.5% in any 1000 Genomes population [51] were downloaded from https://console.cloud. google.com/storage/browser/dm-enformer/variant-scores. The authors do not recommend any specific way of computing a single deleteriousness score. We scored variants using minus the norm of the 5 313 delta features (SNP Activity Difference or SAD). We found that the *L*^1^ norm works better than the *L*^2^ or *L^∞^* norm.
- pLI [33]: precomputed scores for the genes were downloaded from gnomAD v2 and v4 releases [23] under the Constraint sections.
- Heterozygous selection (*hs*) coefficients [34]: precomputed log10 maximum a posteriori (MAP) estimates for the genes were obtained from the supplementary data in the original publication.

### GPN-MSA Captures Variant Functional Impact

A variant’s impact on loss of fitness is mediated by genetic and functional pathways. To investigate whether GPN-MSA captures any functional impact of a variant, we performed functional enrichment analysis on the same random subset of 10M simulated variants in chromosome 22 (see **VEP glossary**). We used 18 functional annotations obtained from the FAVOR database [52] (accessed via Harvard Dataverse on April 10, 2023), which measure both impact of natural selection and gene regulatory activity of a variant (see Supplementary Table 6). For clarity, we collect computational details of the functional annotations and summarize them below.

- B Statistic [53], nucleotide diversity [54] and recombination rate [54] are mathematical quantities derived from evolutionary models, and are computed directly on the genomic position of the variant. They provide population-genetic interpretation of the impact of natural selection on the variant.
- Epigenetic tracks, RNA-seq, DNAse-seq, percent GC and percent CpG were all computed on genomic positions, to be included as training features in CADD [24]. Specifically, ENCODE track features are not gene-specific but are distributed as “bigWig” value tracks along genomic coordinates. Values for each cell-type for which a track is available are summarized to create a new genome coordinate based track, which is subsequently assigned to the variant based on its genomic position. Whenever a variant is not annotatable for a track (e.g., RNA-seq level for a non-exonic variant), an NA value is assigned.

We note that across all functional annotations, at least 69% of all variants (*n* = 6, 854, 981) were annotated. The average completeness rate across all functional annotations was 85%. This ensured that sample sizes were sufficiently large for statistical analyses to be well-powered.

To investigate whether extreme values of GPN-MSA were associated with functional impact, we ran Mann-Whitney tests between the lowest (most deleterious) 1% GPN-MSA scoring (“target”) variants and the remaining (“background”) variants, across all 18 annotations. We found significant enrichment (*p <* 0.05 after controlling for FWER) of 9 histone mark levels, as well as RNA-seq and DNase-seq levels in each dataset, and significant depletion of nucleotide diversity, recombination rate, and B statistic (Supplementary Figure 2). Significant depletions were observed of H3K9me3 and H3K27me3, two recognized gene repressors. These results suggest that extremely negative GPN-MSA scores could potentially prioritize variants with impact on gene expression and regulation.

### Code Availability

Code to reproduce all results is available at https://github.com/songlab-cal/gpn.

### Model and Data Availability

The pretrained model, training dataset, benchmark datasets, and pre-computed scores for all 9 billion possible single nucleotide variants in the human genome are available at https://huggingface. co/collections/songlab/gpn-msa-65319280c93c85e11c803887. Sequence logos derived from GPN-MSA’s predictions can be visualized at https://genome.ucsc.edu/s/gbenegas/gpn-msa-sapiens.

## Acknowledgements

We thank Dr. Martin Kircher for helpful correspondence regarding CADD. This research is supported in part by an NIH grant R35-GM134922 and a grant from the Koret-UC Berkeley-Tel Aviv University Initiative in Computational Biology and Bioinformatics.

## Supplementary Tables

**Supplementary Table 1:** Held-out accuracy. 39M positions in chromosome 22 were evaluated.

| Region | Accuracy | Median<br>perplexity |
| --- | --- | --- |
| CDS | 0.915 | 1.032 |
| 5' UTR | 0.837 | 1.646 |
| 3' UTR | 0.821 | 1.688 |
| Non-exonic | 0.690 | 1.978 |

**Supplementary Table 2:** Held-out accuracy at sites where ancestral allele differs from reference allele. 245K positions in chromosome 22 were evaluated.

| Region | Accuracy<br>predicting<br>reference | Accuracy<br>predicting<br>ancestral |
| --- | --- | --- |
| CDS | 0.227 | 0.680 |
| 5' UTR | 0.177 | 0.679 |
| 3' UTR | 0.245 | 0.603 |
| Non-exonic | 0.268 | 0.515 |

**Supplementary Table 3:**
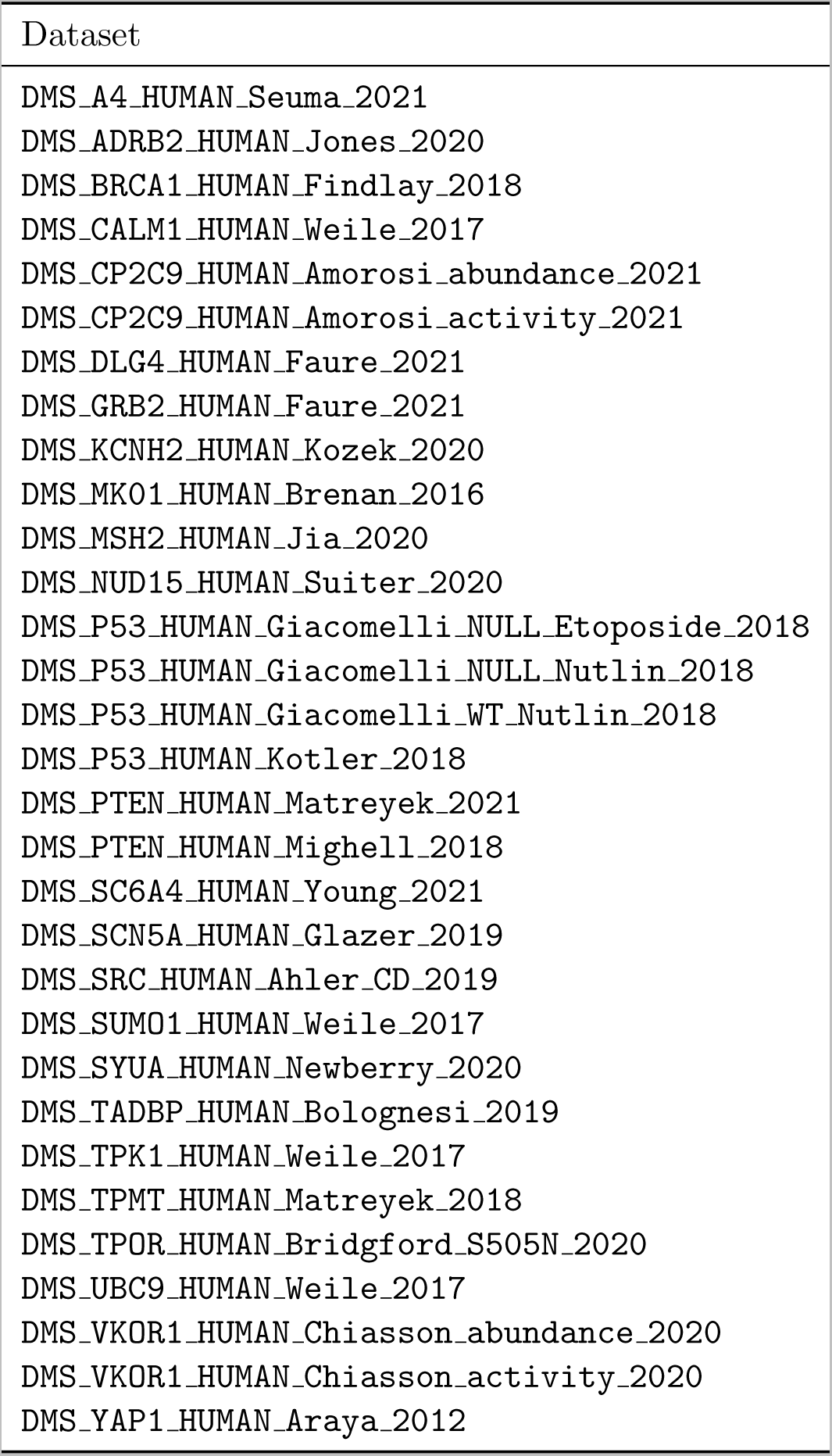
DMS datasets from ProteinGym.

**Supplementary Table 4:**
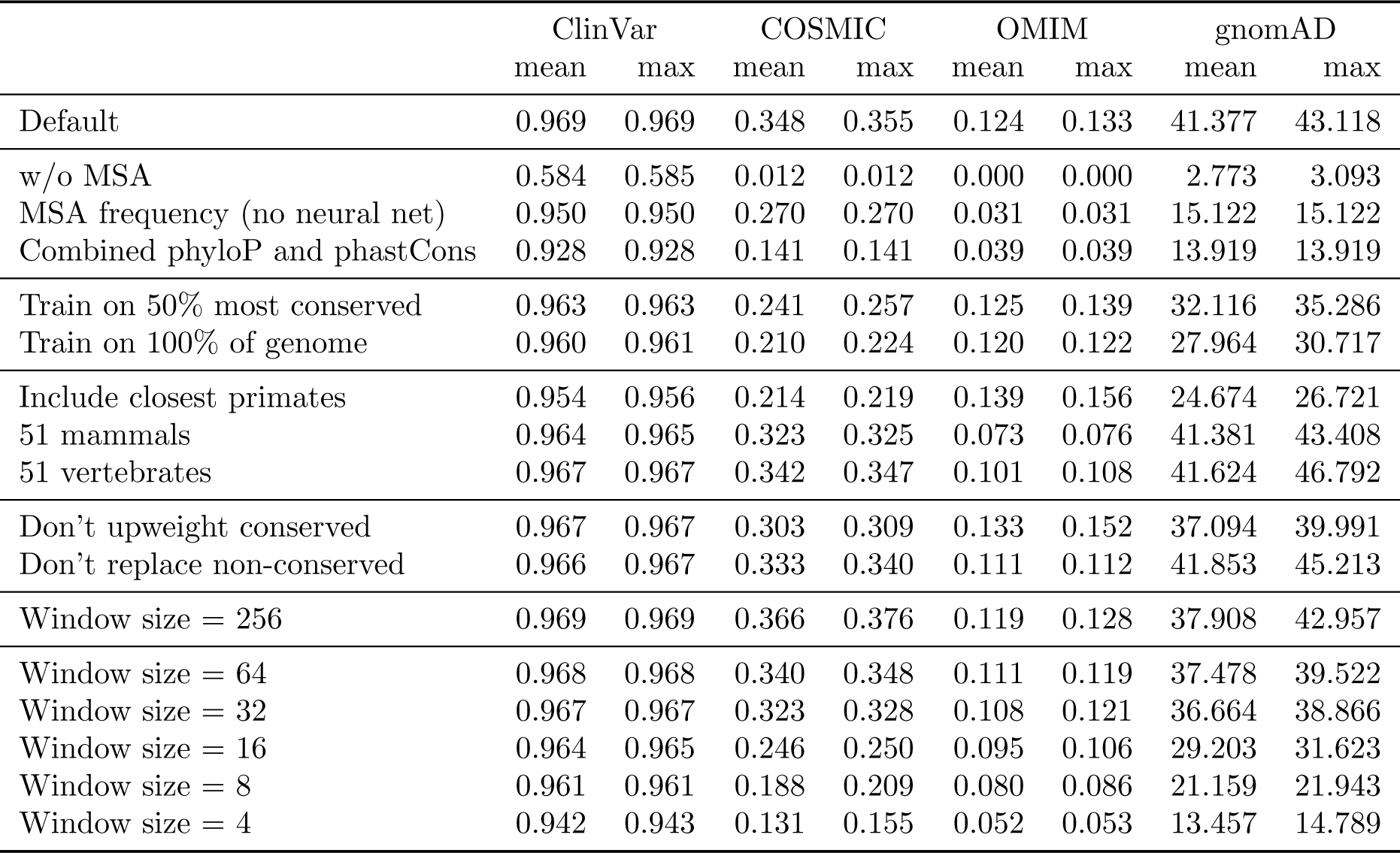
Ablation study. Performance of three random seeds of each independent ablation on four variant effect prediction metrics. ClinVar (AUROC): same setting as Figure 2a. COSMIC (AUPRC): same setting as Figure 2b. OMIM (AUPRC): same setting as Figure 2c. gnomAD (odds ratio): same approach as Figure 2d, including all variant types, but subsetting the rare variants to match the number of common variants. Ablations are detailed in Methods.

|  | ClinVar |  | COSMIC |  | OMIM |  | gnomAD |  |
| --- | --- | --- | --- | --- | --- | --- | --- | --- |
|  | mean | max | mean | max | mean | max | mean | max |
| Default | 0.969 | 0.969 | 0.348 | 0.355 | 0.124 | 0.133 | 41.377 | 43.118 |
| w/o MSA | 0.584 | 0.585 | 0.012 | 0.012 | 0.000 | 0.000 | 2.773 | 3.093 |
| MSA frequency (no neural net) | 0.950 | 0.950 | 0.270 | 0.270 | 0.031 | 0.031 | 15.122 | 15.122 |
| Combined phyloP and phastCons | 0.928 | 0.928 | 0.141 | 0.141 | 0.039 | 0.039 | 13.919 | 13.919 |
| Train on 50% most conserved | 0.963 | 0.963 | 0.241 | 0.257 | 0.125 | 0.139 | 32.116 | 35.286 |
| Train on 100% of genome | 0.960 | 0.961 | 0.210 | 0.224 | 0.120 | 0.122 | 27.964 | 30.717 |
| Include closest primates | 0.954 | 0.956 | 0.214 | 0.219 | 0.139 | 0.156 | 24.674 | 26.721 |
| 51 mammals | 0.964 | 0.965 | 0.323 | 0.325 | 0.073 | 0.076 | 41.381 | 43.408 |
| 51 vertebrates | 0.967 | 0.967 | 0.342 | 0.347 | 0.101 | 0.108 | 41.624 | 46.792 |
| Don't upweight conserved | 0.967 | 0.967 | 0.303 | 0.309 | 0.133 | 0.152 | 37.094 | 39.991 |
| Don't replace non-conserved | 0.966 | 0.967 | 0.333 | 0.340 | 0.111 | 0.112 | 41.853 | 45.213 |
| Window size = 256 | 0.969 | 0.969 | 0.366 | 0.376 | 0.119 | 0.128 | 37.908 | 42.957 |
| Window size = 64 | 0.968 | 0.968 | 0.340 | 0.348 | 0.111 | 0.119 | 37.478 | 39.522 |
| Window size = 32 | 0.967 | 0.967 | 0.323 | 0.328 | 0.108 | 0.121 | 36.664 | 38.866 |
| Window size = 16 | 0.964 | 0.965 | 0.246 | 0.250 | 0.095 | 0.106 | 29.203 | 31.623 |
| Window size = 8 | 0.961 | 0.961 | 0.188 | 0.209 | 0.080 | 0.086 | 21.159 | 21.943 |
| Window size = 4 | 0.942 | 0.943 | 0.131 | 0.155 | 0.052 | 0.053 | 13.457 | 14.789 |

**Supplementary Table 5:** Training hyperparameters.

|  |  |
| --- | --- |
| Weight decay | 0.01 |
| Max steps | 30 K |
| Batch size | 2048 |
| Learning rate | $10^{-4}$ |
| Learning rate schedule | Cosine |
| Learning rate warmup steps | 1 K |

**Supplementary Table 6:**
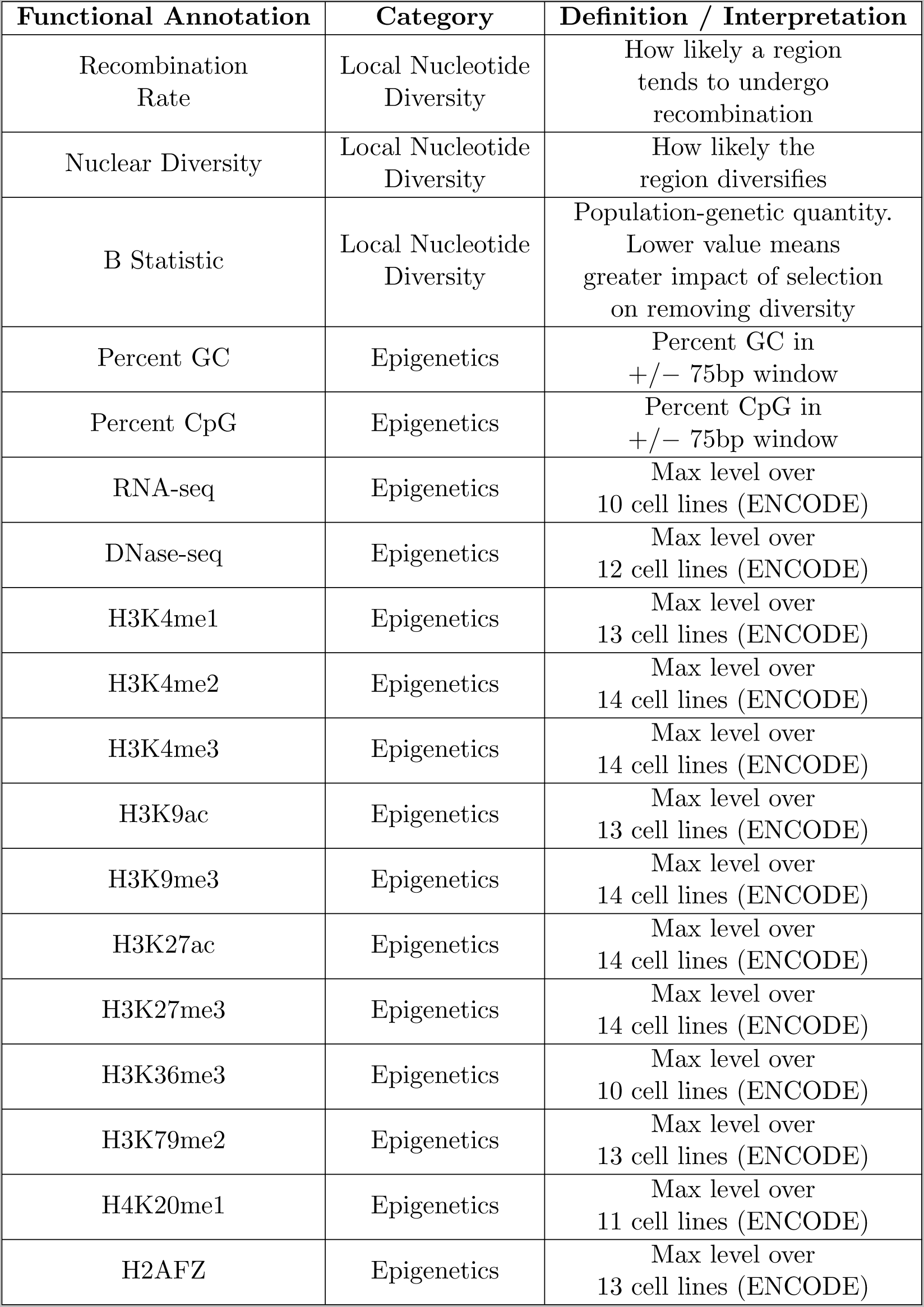
Functional annotations considered in our analysis of functional enrichment. In particular, annotations relying on predictive models are not considered.

## Supplementary Figures

**Supplementary Figure 1:**
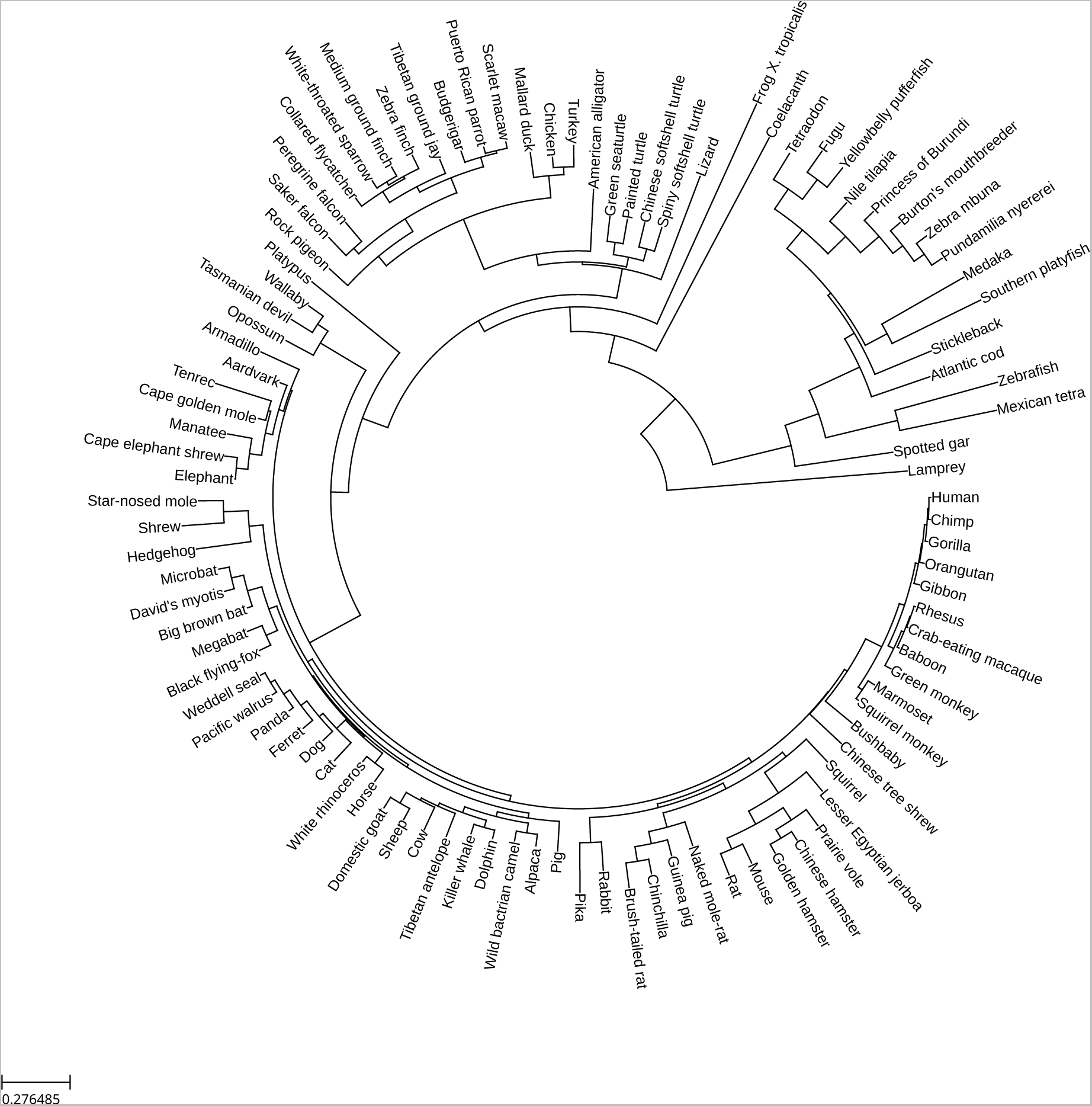
Phylogenetic tree of 100 vertebrates.

**Supplementary Figure 2:**
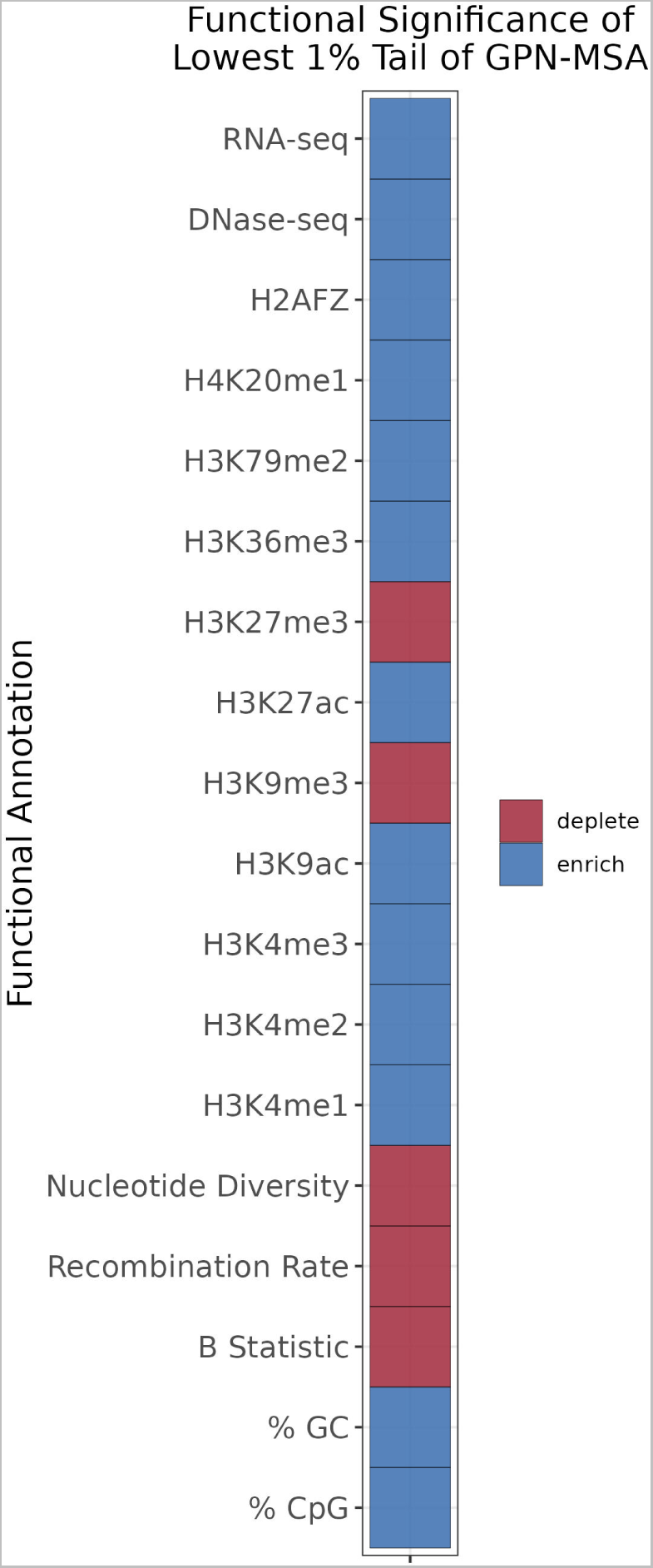
Functional enrichment and depletion of deleterious tail. Significant (Wilcoxon-Mann-Whitney test with FWER controlled at 0.05) enrichments and depletions of functional annotations between the deleterious GPN-MSA tail set of variants and background variants. For tracks like RNA-seq, which require the variant to be exonic, variants without the annotation are not included in the two-sample test.

**Supplementary Figure 3:**
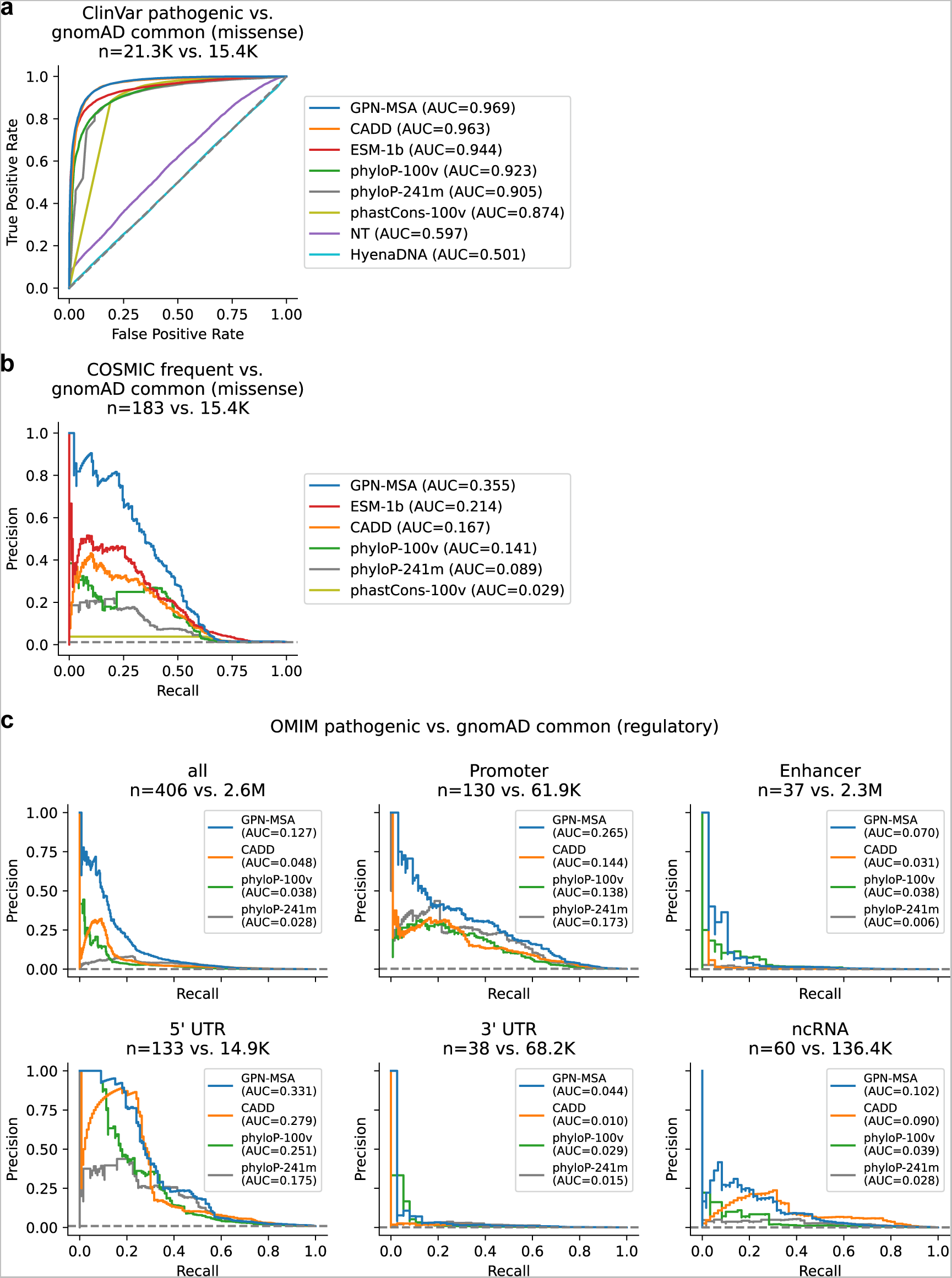
Receiver Operating Characteristic and Precision-Recall curves for variant effect prediction. (a) Same setting as Figure 2a. **(b)** Same setting as Figure 2b. **(c)** Same setting as Figure 2c.

**Supplementary Figure 4:**
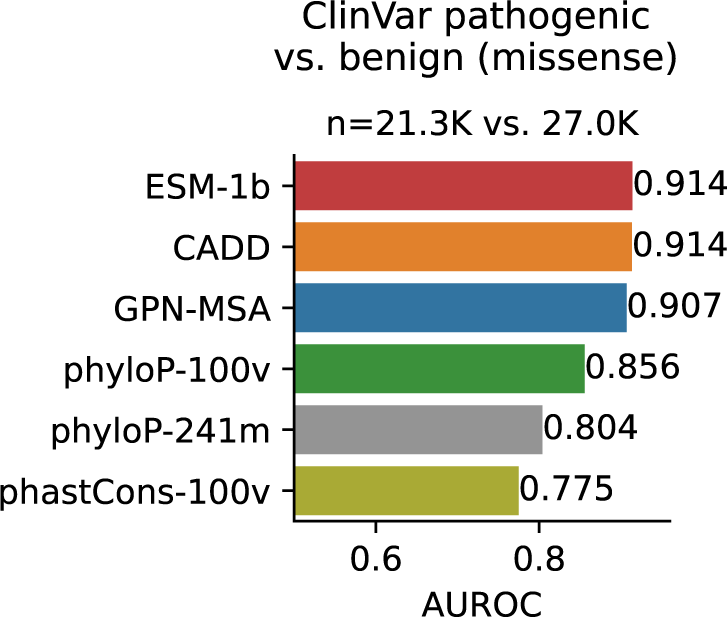
ClinVar pathogenic vs. benign.

**Supplementary Figure 5:**
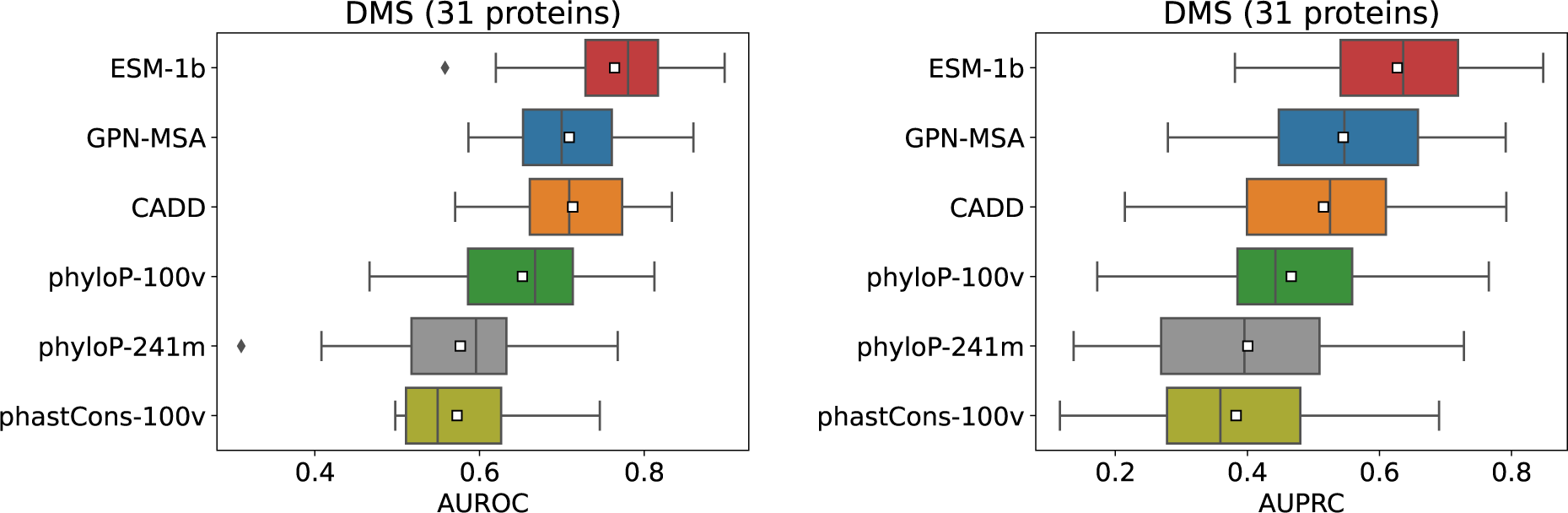
Performance on 31 protein DMS data sets from ProteinGym. AUROC and AUPRC results for binary classification (Methods) are summarized.

**Supplementary Figure 6:**
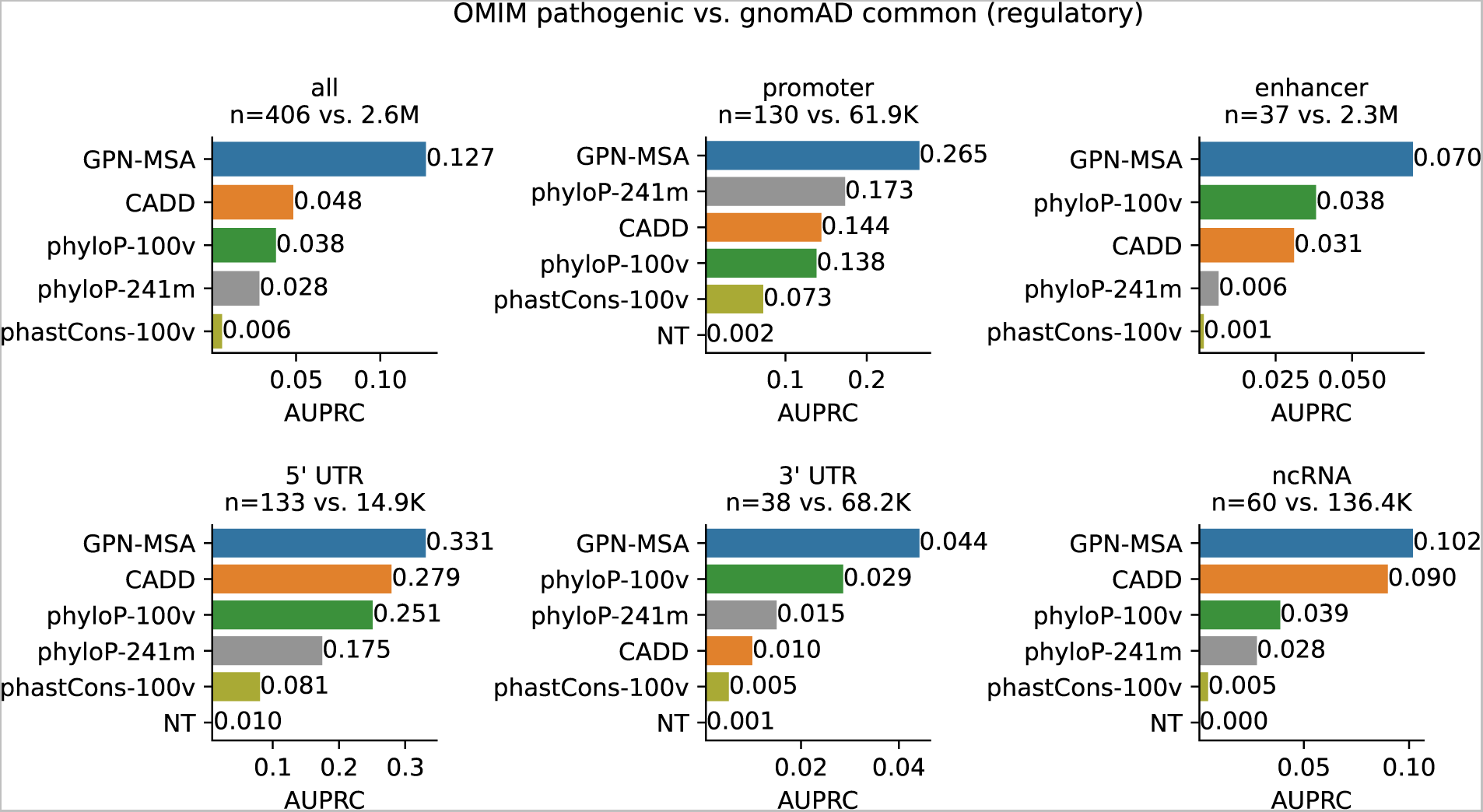
OMIM performance for specific variant categories.

**Supplementary Figure 7:**
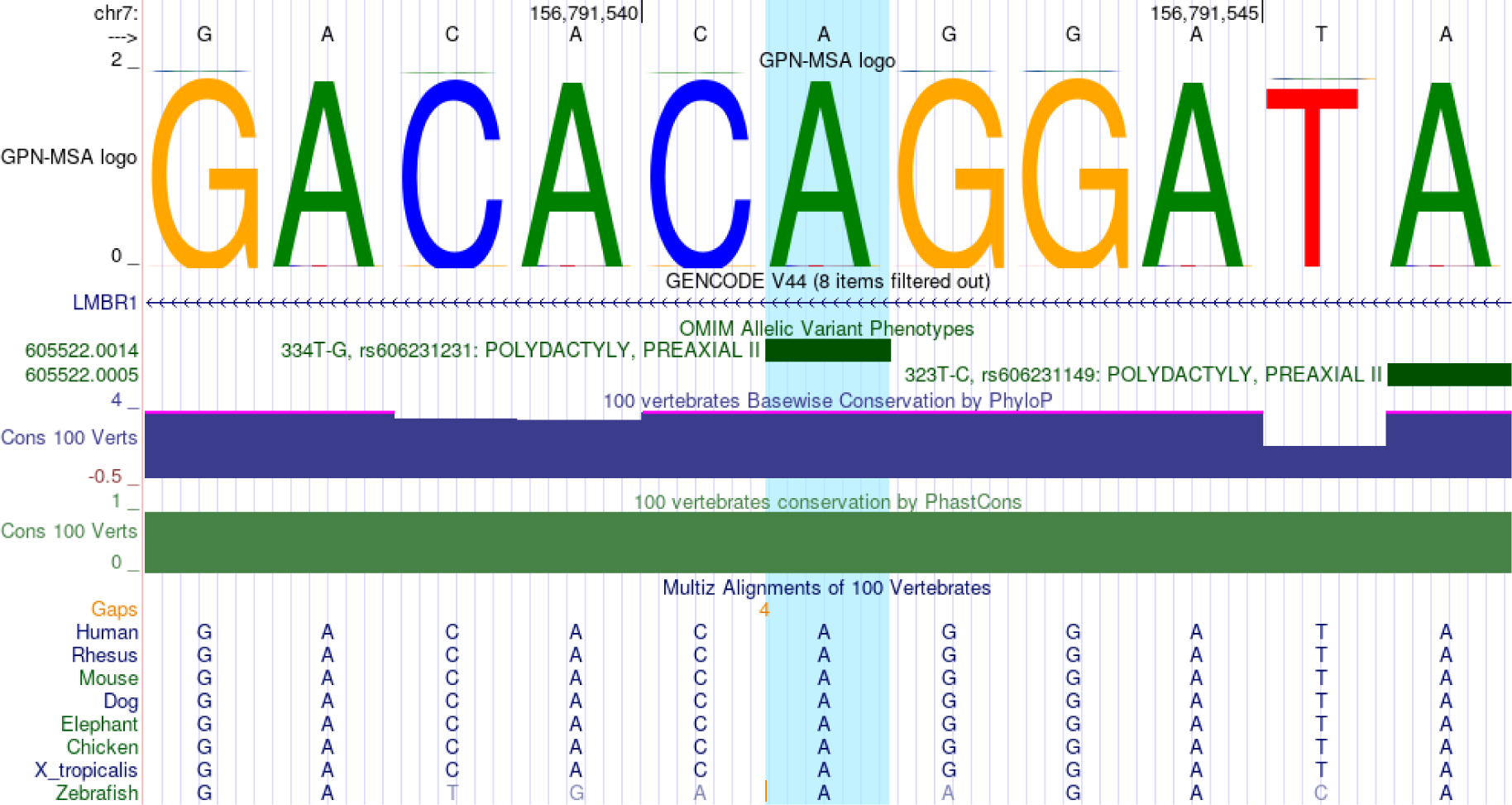
Example enhancer variant: rs606231231. The variant is ranked at the following percentiles: GPN-MSA: 0.0005%, phyloP-100v: 0.0008%, phyloP-241m: 0.3%, CADD: 0.3%, phastCons-100v: 0.3%.

**Supplementary Figure 8:**
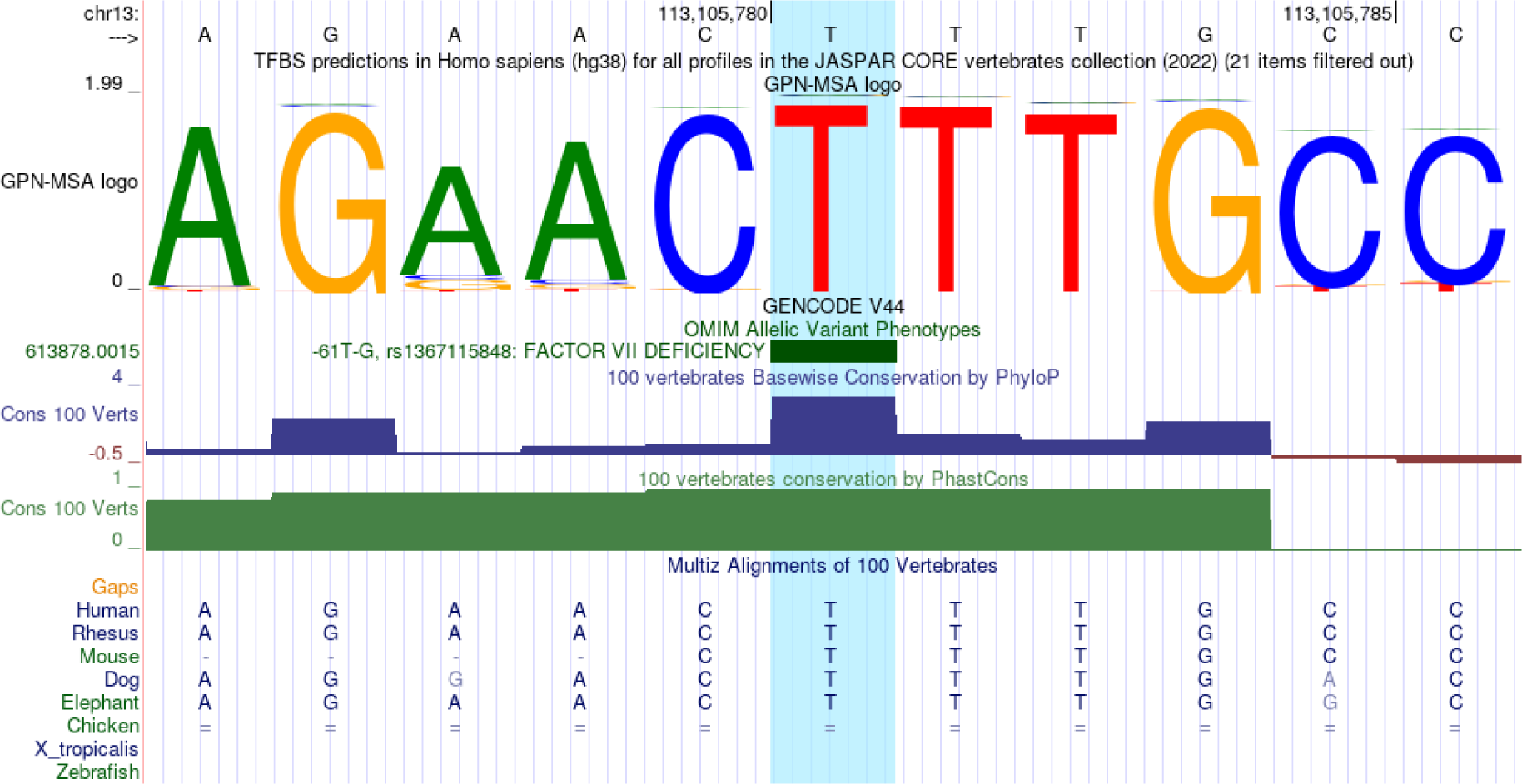
Example promoter variant: rs1367115848. The variant is ranked at the following percentiles: GPN-MSA: 0.01%, phyloP-100v: 0.3%, phastCons-100v: 0.5%, CADD: 0.8%, phyloP-241m: 0.9%, NT: 30.0%.

**Supplementary Figure 9:**
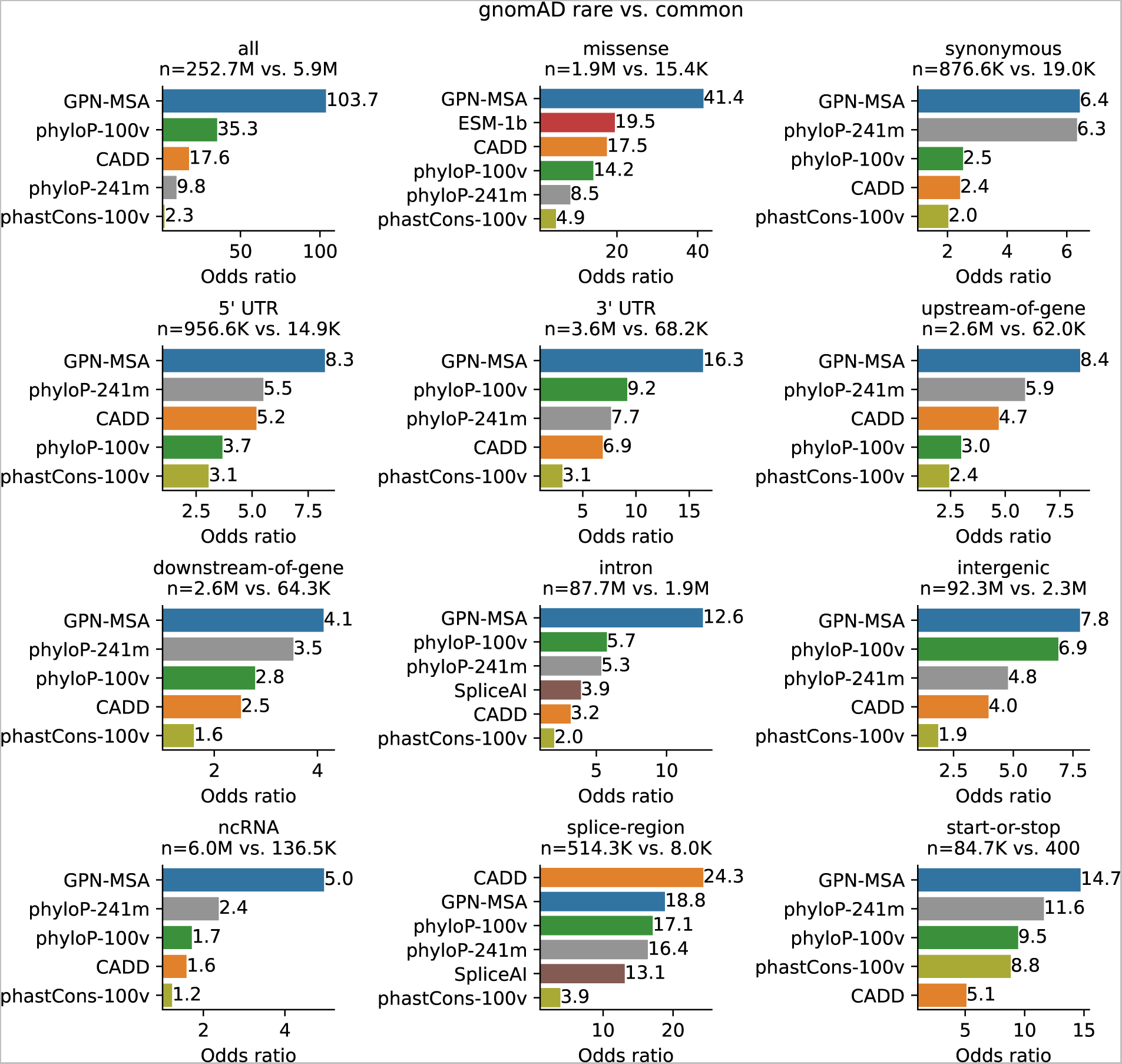
gnomAD rare vs. common odds ratios for different variant categories. Same setting as Figure 2f. Models with non-significant *p*-values are not displayed.

**Supplementary Figure 10:**
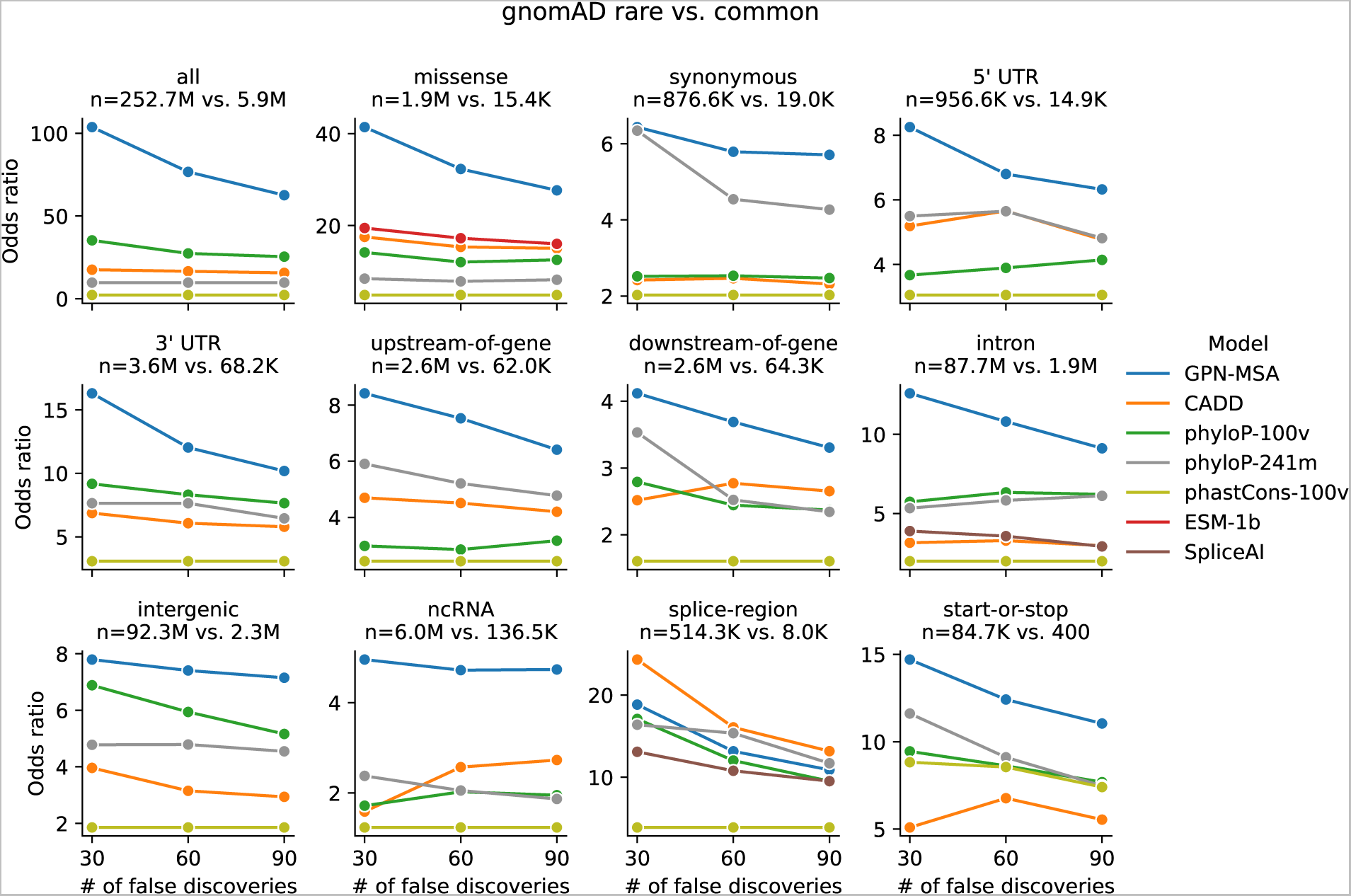
gnomAD rare vs. common odds ratios varying the number of false discoveries. Same setting as Figure 2f. Models with non-significant *p*-values are not displayed.

**Supplementary Figure 11:**
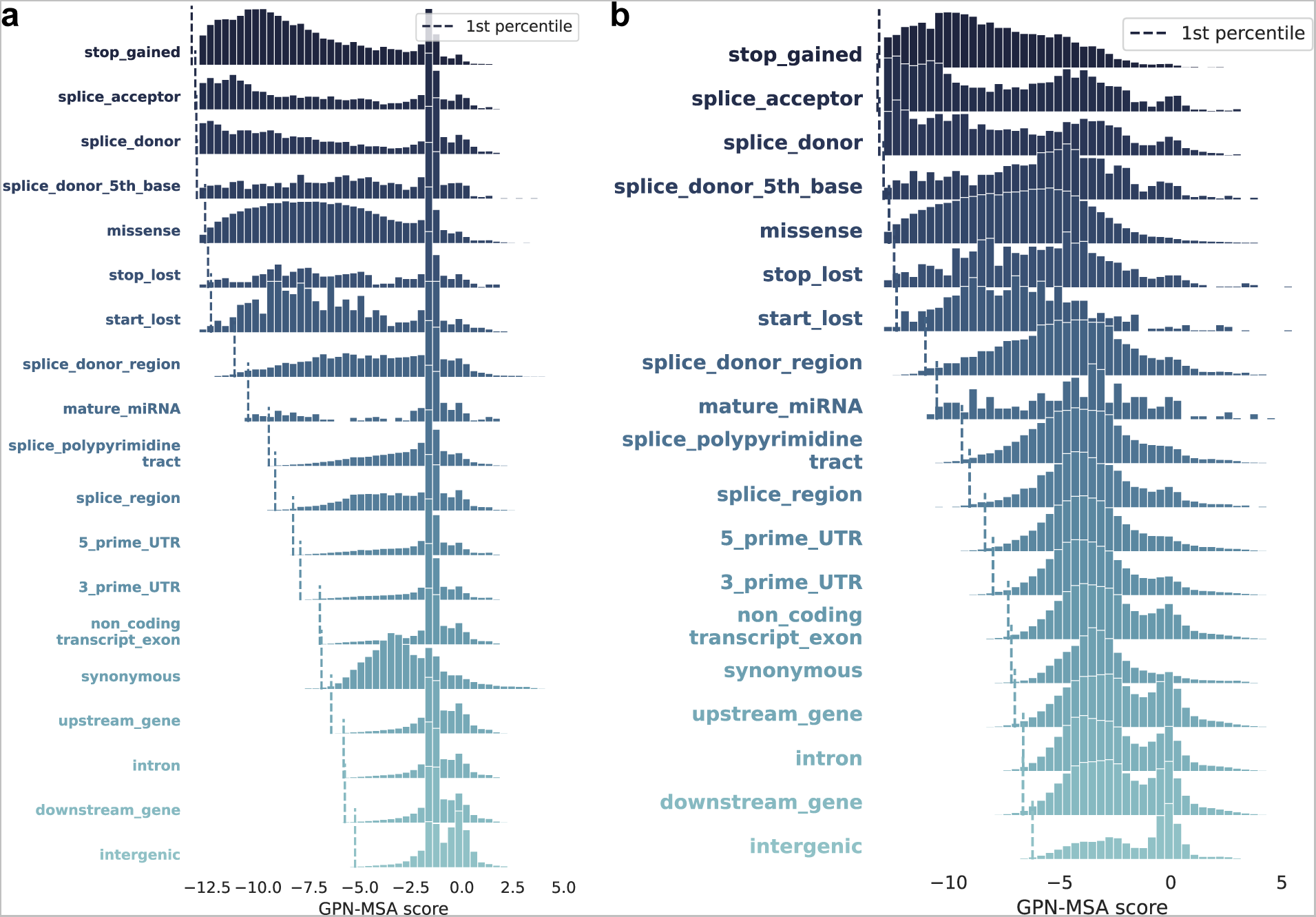
*In silico* mutagenesis. Distribution of GPN-MSA scores of a random subset of 10M SNPs in held-out chromosome 22, across categories, sorted by first percentile (dashed vertical lines). **(a)** Default model. **(b)** Without random replacement with another nucleotide at non-conserved positions.

**Supplementary Figure 12:**
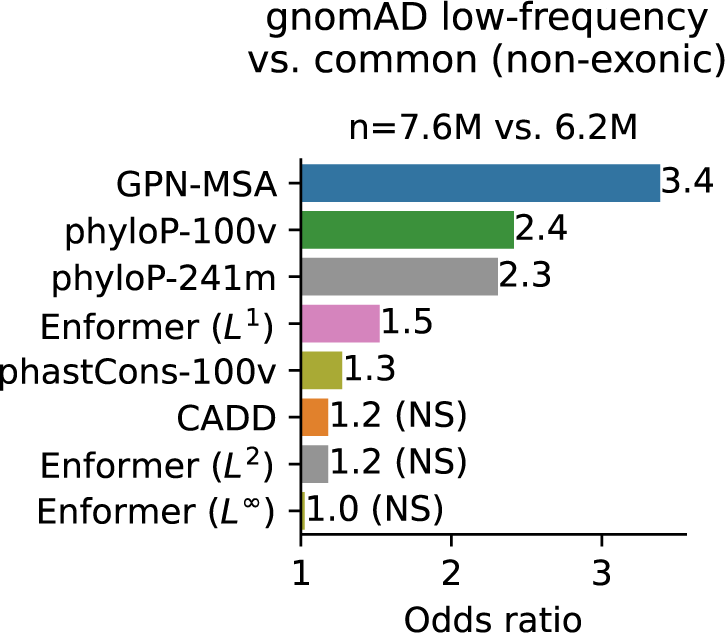
Comparison with Enformer. Enrichment of low-frequency (0.5% *<* AF *<* 5%) vs. common (MAF *>* 5%) gnomAD non-exonic variants in the tail of deleterious scores (the threshold was chosen such that each score makes 30 false discoveries).

**Supplementary Figure 13:**
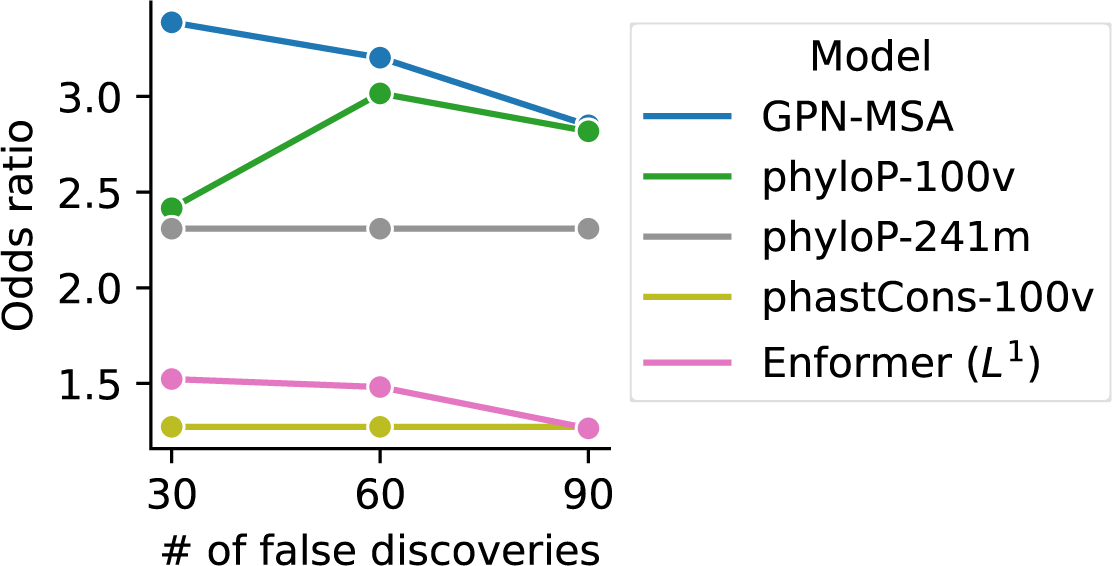
gnomAD low-frequency vs. common odds ratios varying number of false discoveries. Same setting as Supplementary Figure 12. Models with non-significant *p*-values are not displayed.

**Supplementary Figure 14:**
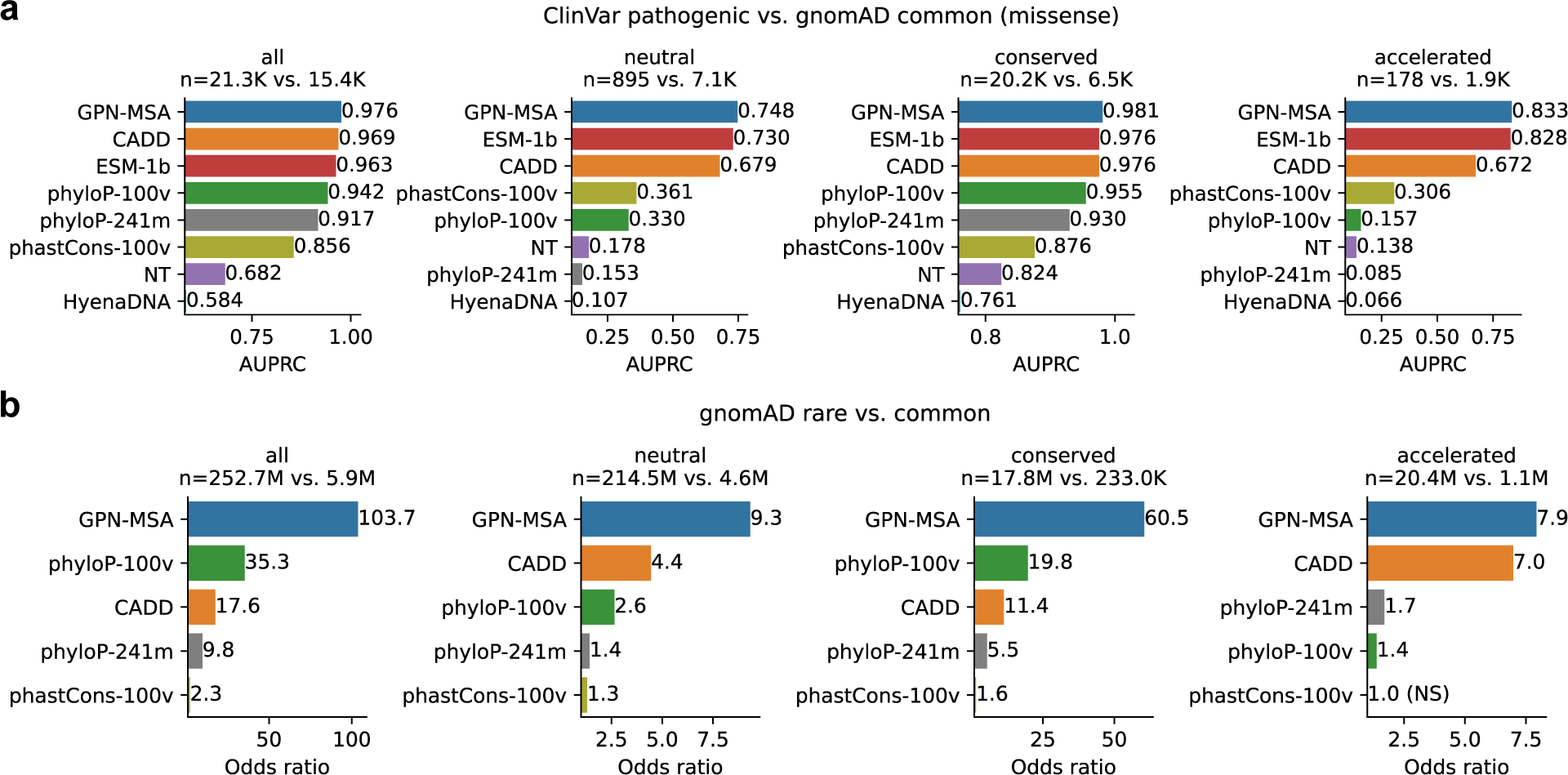
Performance stratified by putative evolutionary class. Conservation and acceleration were defined using phyloP-241m cutoffs at *p <* 0.05. **(a)** Same setting as Figure 2c, using AUPRC instead of AUROC since the class balance varies substantially. **(b)** Same setting as Figure 2f.

**Supplementary Figure 15:**
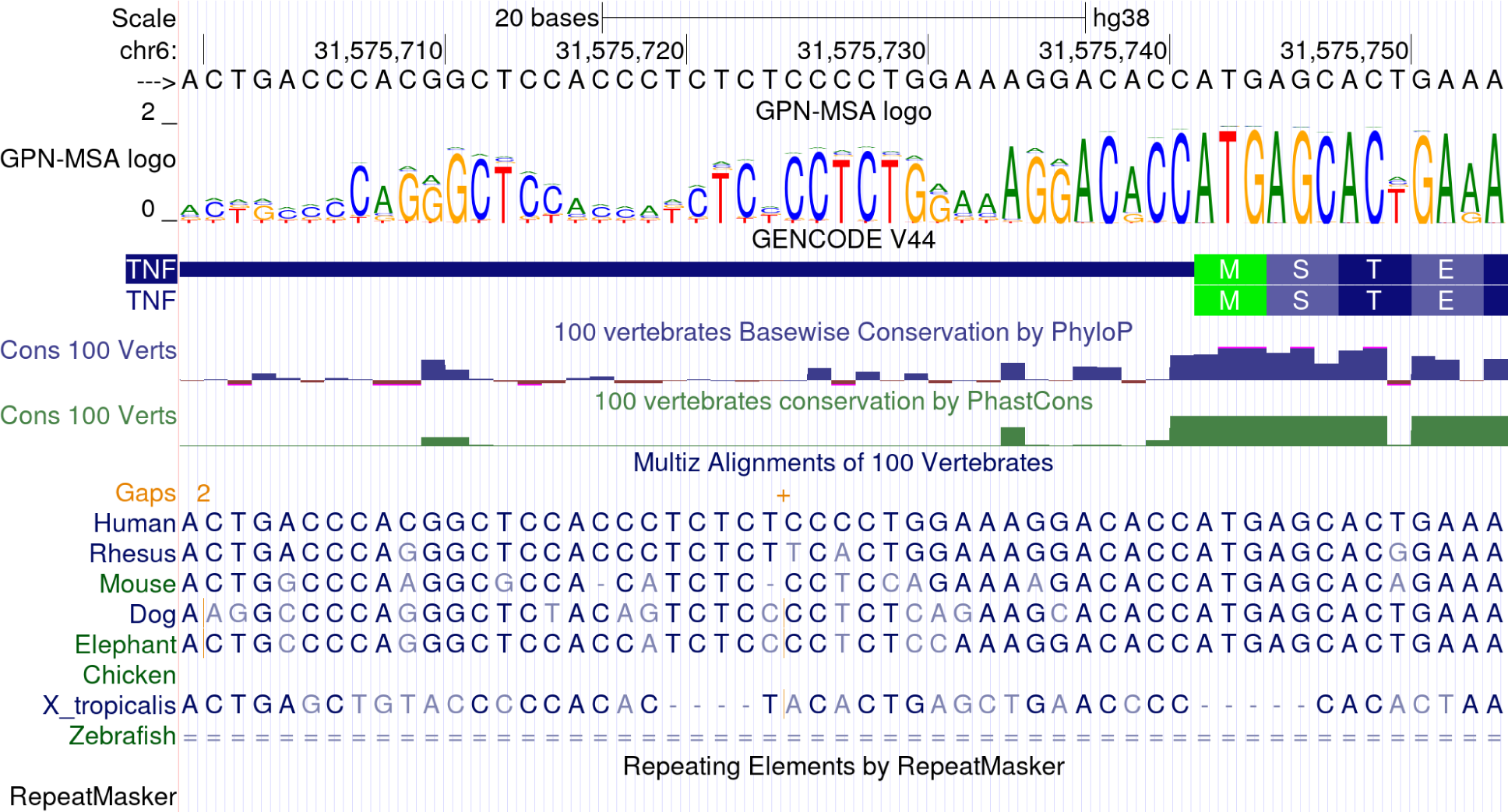
GPN-MSA logo track on the UCSC Genome Browser. Shown region: chr6:31,575,700-31,575,754.

**Supplementary Figure 16:**
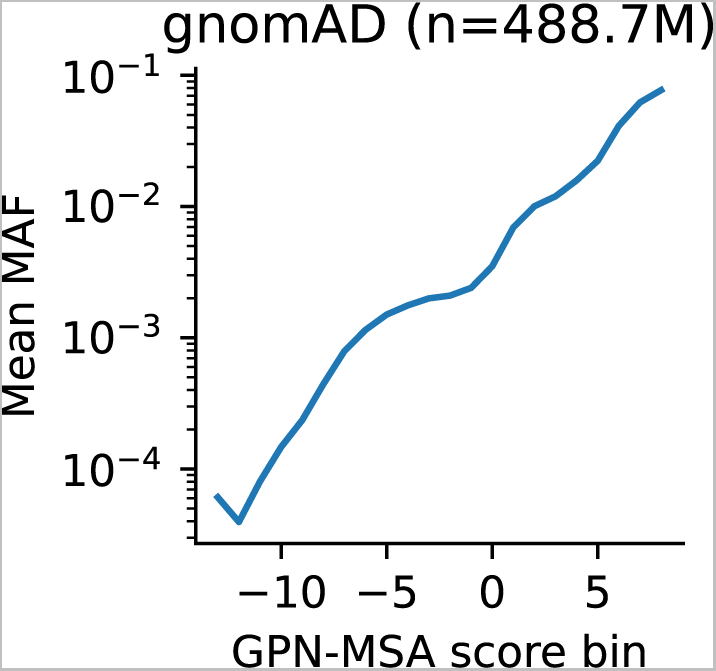
Mean minor allele frequency (MAF) for different bins of GPN-MSA scores ([*−*13.5*, −*12.5), [*−*12.5*, −*11.5)*, …,* [8.5, 9.5)) in the full set of gnomAD bi-allelic sites.

**Supplementary Figure 17:**
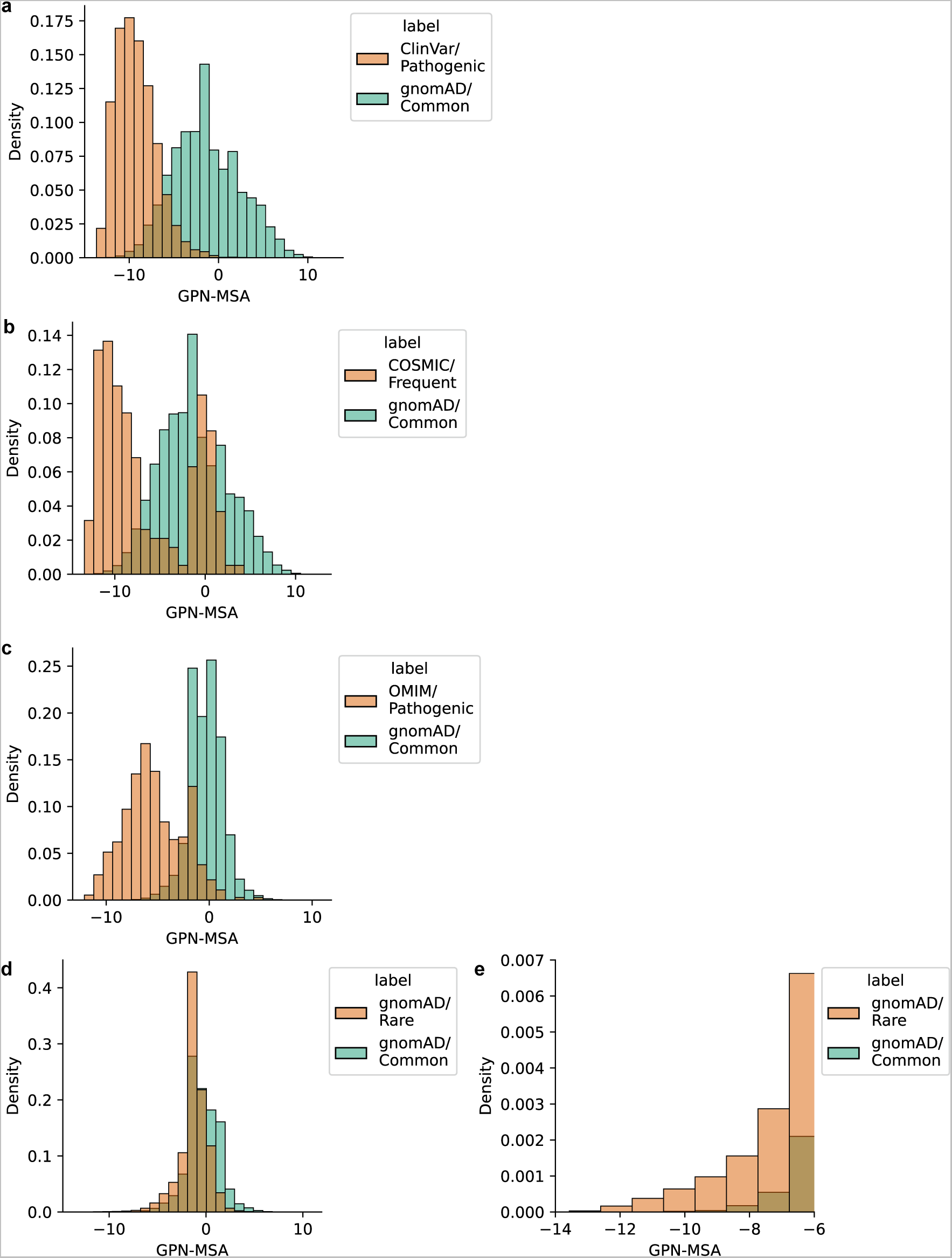
Histogram of GPN-MSA scores. **(a)** Scores for Figure 2c. **(b)** Scores for Figure 2d. **(c)** Scores for Figure 2e. **(d)** Scores for Figure 2f. **(e)** A zoomed-in version of (d) highlighting the left tail.

## Footnotes

1 https://huggingface.co/docs/transformers/model_doc/roformer#transformers.RoFormerConfig

